# Early Postnatal Dysfunction of ACC PV Interneurons in Shank3B^-/-^ Mice

**DOI:** 10.1101/2024.10.04.616584

**Authors:** Yi-Chun Shih, Lars Nelson, Michael Janeček, Michael Matarazzo, Andrew D’Agostino, Rui T. Peixoto

**Affiliations:** Department of Psychiatry, University of Pittsburgh, 450 Technology Dr, Pittsburgh, PA 15219, USA; Center for Neuroscience at the University of Pittsburgh

## Abstract

Anterior cingulate cortex **(ACC)** dysfunction is implicated in the cognitive and social deficits associated with autism spectrum disorder **(ASD)**, yet the developmental trajectory of ACC circuit maturation in ASD remains poorly understood. Here, we examined the postnatal development of glutamatergic synaptic connectivity and intrinsic excitability in layer 2/3 pyramidal neurons **(PYR)** and Parvalbumin-expressing interneurons **(PVIN)** in the ACC of mice harboring a deletion in SHANK3 **(Shank3B^-/-^)**, a well-established genetic cause of autism. We found that ACC PVINs in Shank3B^-/-^ mice exhibit reduced excitability and *in vivo* hypoactivity as early as postnatal day 15 **(P15)** despite receiving normal levels of glutamatergic input. This early PVIN hypoexcitability is associated with decreased feedforward inhibition from the mediodorsal thalamus and reduced hyperpolarization-activated **(I_h_)** currents mediated by hyperpolarization-activated cyclic nucleotide gated **(HCN)** channels. In contrast, PYRs display normal excitability and synaptic input at this stage but already exhibit reduced I_h_ currents, indicating an early emergence of HCN channel dysfunction in both PYR and PVIN. By adulthood, both neuron populations undergo marked phenotypic changes, characterized by reduced glutamatergic synaptic input and divergent alterations in excitability. Together, these findings reveal a distinct sequence of early PVIN dysfunction followed by cell-type specific circuit reorganization within ACC layer 2/3 of Shank3B^-/-^ mice and identify HCN channelopathy and impaired PVIN-mediated inhibition as early pathogenic features of SHANK3-related neurodevelopmental disorders.

## Introduction

Autism spectrum disorders **(ASD)** are marked by significant cognitive deficits that typically emerge in the first years of life^1^, implicating abnormal maturation of the medial prefrontal cortex **(mPFC)** as a central factor in ASD pathogenesis^2^. Neuroimaging studies have consistently reported cortical hypertrophy and disrupted functional connectivity in mPFC regions of young children with ASD^2–7^. In parallel, epilepsy and subclinical epileptiform activity are highly comorbid with ASD^8–11^, pointing to dysregulated cortical excitability. Mouse models carrying ASD-associated mutations further support these observations and frequently exhibit altered neuron function and abnormal patterns of activity in frontal cortical regions^12–15^. However, the vast majority of these studies have focused on adult animals, leaving a critical gap in our understanding of how the development of frontal cortical circuits is affected in these conditions^16^. Despite the essential role of the mPFC in early cognitive and social development^17^, the specific mechanisms underlying the early emergence of prefrontal circuit dysfunction in ASD remain poorly understood.

The high prevalence of seizures and cortical activity imbalances in individuals with autism suggests that deficits in inhibitory mechanisms are a common pathophysiological feature of ASD^18–20^. In particular, alterations in GABAergic Parvalbumin **(PV)** expressing interneurons **(PVIN)** have been widely observed in transcriptomic^21,22^ and postmortem histological studies of individuals with autism^23–25^. Similarly, multiple mouse models carrying ASD-linked mutations exhibit disrupted PVIN function, abnormal cortical PV expression patterns^26–32^ and impaired synaptic connectivity^33,34^. PVINs are critical regulators of cortical circuit activity, providing tightly controlled inhibition to local pyramidal neurons **(PYR)**^35,36^. This fast and temporally precise inhibition is supported by specialized synaptic and intrinsic membrane properties^37^. Notably, several ASD risk genes converge on pathways that govern neuronal excitability, glutamatergic synapse development, and GABAergic signaling^20,38–40^, highlighting the postnatal maturation of PVIN as a potential locus of vulnerability in ASD. Given their critical role in mediating cortical inhibition, disruptions in PVIN development may imbalance early cortical activity, impairing critical neurodevelopmental processes that rely on properly timed neural activity. However, studying the early postnatal maturation of cortical PVINs has been hindered by the reliance on PV expression for cell identification, a protein that in mice is not expressed in frontal cortical areas until the third postnatal week^41^. Consequently, the mechanisms regulating early PVIN maturation in frontal cortical circuits remain poorly understood.

Mutations in SH3 and Multiple Ankyrin Repeat Domains Protein 3 **(SHANK3)** are highly penetrant causes of autism^39,40^ and are associated with epilepsy and profound cognitive impairments^9,42,43^. SHANK3 is a complex gene that encodes a family of postsynaptic scaffolding proteins that regulate the maturation and function of glutamatergic synapses^44^. Deletion of exons Δ13-16 encoding the PDZ domain of SHANK3 in mice **(Shank3B^-/-^)** alters neuronal excitability^45–49^ and causes behavioral deficits and hypoconnectivity of frontal cortical circuits^50–53^. Notably, adult Shank3B^-/-^ mice show reduced activity of GABAergic interneurons and increased PYR activity in primary somatosensory cortex in response to sensory stimulation^26,54^. Altered cortical expression of PV is also one of the most replicated findings in Shank3B^-/-^ mice^26,27,32^, strongly implicating PVIN dysfunction. However, the specific effects of loss of Shank3 on PVIN function have not been characterized.

Here we investigated how the loss of Shank3 disrupts the postnatal maturation of layer 2/3 **(L2/3)** circuits in the anterior cingulate cortex **(ACC)** by examining the developmental trajectory of glutamatergic synaptic connectivity, membrane excitability and *in vivo* activity patterns of PYR and PVIN in Shank3B^-/-^ mice. We focused on the ACC because structural and functional deficits in this frontal cortical region are strongly linked to autism^56–59^, and deficits in ACC PYR synaptic connectivity and function have been reported in adult Shank3B^-/-^ mice^53^. Our findings reveal that L2/3 ACC PVIN exhibit significant hypoexcitability as early as postnatal day **(P15)**. In addition, PVIN show reduced inhibitory output onto PYR and a substantial reduction in feedforward inhibition **(FFI)** of thalamocortical inputs. At this stage, L2/3 ACC PYR display normal synaptic and intrinsic properties, but both PYR and PVIN exhibit reduced hyperpolarization-activated cation channel **(HCN)** currents, consistent with early onset of HCN deficits in both cell types. *In vivo* two-photon calcium imaging confirmed altered patterns of cortical activity in the ACC of postnatal Shank3B^-/-^ mice, showing less frequent calcium transients in PVIN. By adulthood, we observed extensive developmental adaptations in L2/3 ACC circuits, with PVIN becoming hyperexcitable and PYR neurons exhibiting hypoexcitability, along with a marked reduction in glutamatergic synaptic inputs in both neuron types. These findings indicate that early postnatal PVIN hypoexcitability, along with HCN channelopathy in both PVINs and PYR are early functional abnormalities in L2/3 ACC circuits of Shank3B^-/-^ mice. Additionally, our results highlight the importance of understanding the developmental trajectory of neural circuit maturation in ASD to distinguish between primary deficits and secondary developmental adaptations.

## Results

### Normal excitability and AMPAR-mediated synaptic inputs of ACC L2/3 PYR in P15 Shank3B^-/-^ mice

Adult Shank3B^-/-^ mice exhibit reduced glutamatergic input in ACC L2/3 PYR^53^. However, many behavioral deficits in these animals are already observed at P15^60,61^, suggesting an early onset of ACC circuit dysfunction. To determine whether loss of Shank3 affects the postnatal maturation of glutamatergic synapses in ACC L2/3 PYR, we recorded AMPAR miniature excitatory postsynaptic currents **(mEPSC)** in acute brain slices of P15 wild-type **(WT)** and Shank3B^-/-^ mice, which provide an estimate of glutamatergic synapse number and strength. Whole-cell recordings in L2/3 PYR were conducted using a cesium-based internal solution in the presence of the voltage-gated sodium channel blocker tetrodotoxin to prevent action potential **(AP)** propagation and evoked synaptic vesicle release with membrane potential **(V_m_)** was clamped at -70mV (Fig. 1). We observed no difference in the amplitude of AMPAR mEPSCs between ACC L2/3 PYR of P15 WT and Shank3B^-/-^ mice (Fig. 1C). In addition, Shank3B^-/-^ PYR showed no difference in average mEPSC frequency compared to WT (Fig. 1D), suggesting a normal number and strength of AMPAR-mediated synapses at P15. No differences in mEPSC rise and decay times, or membrane capacitance were observed (Supplementary Fig. 1A–1C and 1E). These findings indicate that the early postnatal maturation of AMPAR inputs onto ACC L2/3 PYR is largely unaffected by loss of Shank3.

**Figure 1.**
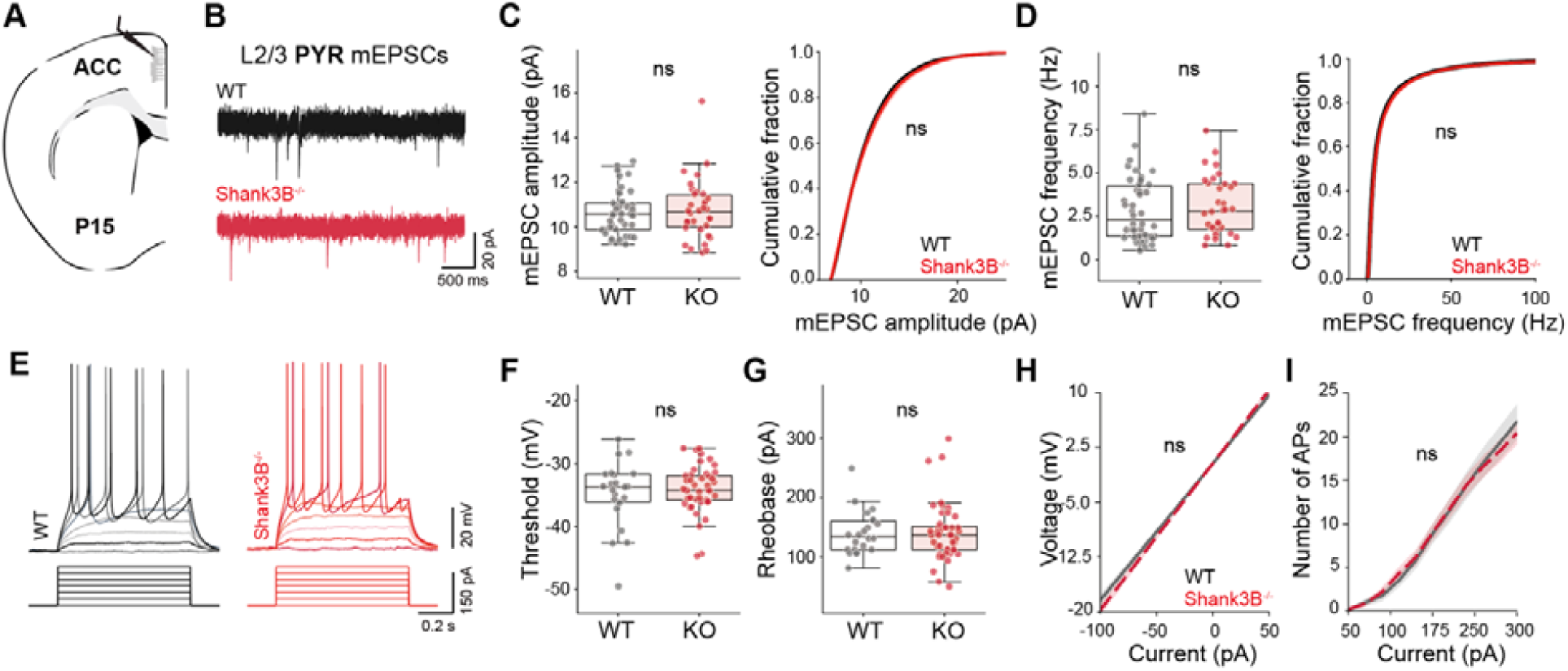
Normal AMPAR-mediated synaptic function and excitability of ACC L2/3 PYR in P15 Shank3B^-/-^ mice. **(A)** Schematic representing coronal brain section and whole-cell recordings of L2/3 PYR in ACC of P15 mice. **(B)** Representative AMPAR mEPSCs in L2/3 ACC PYR of WT (black) and Shank3B^-/-^ (red) mice. **(C)** Average (left) and cumulative distribution (right) of mEPSC amplitudes (Average±SD, WT 10.6±1.0pA; KO 10.7±1.3pA, unpaired Mann Whitney (MW) test p=0.88. Kolmogorov-Smirnov (KS) test p=1.0) and **(D)** frequency per neuron (Average±SD, WT 2.9±1.8Hz; KO 3.1±1.7Hz, MW test p=0.54. KS test p=0.98). **(E)** Representative current-clamp recording depicting action potential firing in response to current injection in L2/3 ACC PYR of WT (black) and Shank3B^-/-^ (red) mice. **(F)** Average AP threshold potential (Average±SD, WT -34.8±5.3mV; KO -34.1±3.7mV, MW test p=0.9) and **(G)** Rheobase current (Average±SD, WT 140.6±37.3pA; KO 141.4±49.2pA, MW test p=0.89). **(H)** Voltage-current (I-V) relationship of L2/3 ACC PYR (Two-way ANOVA I-V p=0.22 for genotype). **(I)** Action potential firing-current (I-F) relationship of L2/3 ACC PYR (Two-way ANOVA I-F p=1.0 for genotype). For mEPSC recording, n=42,35 neurons from n=4,3 WT and Shank3B^-/-^ mice. For current-clamp recordings, n=25,44 neurons from n=3,4 WT and Shank3B^-/-^ mice. Box plots show median and quartile values. eCDFs represent average of CDF from mEPSCs of each recorded neuron. Box plots *P<0.05 with MW test. eCDFs *P<0.05 with KS test. Regression plots *P<0.05 with two-way ANOVA.

Shank3 deficiency is associated with abnormal intrinsic excitability^45–49^. To determine whether the intrinsic properties of ACC L2/3 PYR are altered in P15 Shank3B^-/-^ mice, we performed current clamp whole-cell recordings at 32C with a potassium based internal solution, and measured membrane voltage changes in response to stepped current injections. We detected no difference in AP threshold potential or rheobase current between ACC L2/3 PYR of both genotypes (Fig. 1F and 1G), as well as no difference in the relationship between current-voltage **(I-V)** and current-AP firing rate **(I-F)** (Fig. 1H and 1I). Resting membrane potential **(RMP)** and AP kinetics were also unaffected (Supplementary Fig. 1F–1L), suggesting normal excitability of ACC L2/3 PYR neurons at P15 in Shank3B^-/-^ mice.

## Reduced excitability but normal AMPAR-mediated synaptic inputs of ACC L2/3 PVIN in P15 Shank3B^-/-^ mice

To determine whether loss of Shank3 alters glutamatergic innervation of ACC PVIN during early postnatal development, we recorded AMPAR-mediated mEPSCs in PVIN of L2/3 ACC of P15 WT and Shank3B^-/-^ mice. To label ACC PVIN, we injected P7 mice in ACC with AAV9-S5E2-tdTom that expresses tdTomato under control of the *Scn1a* E2 (S5E2) enhancer element ^62^ (Fig. 2A and 2E). This approach results in enriched labeling of PVIN which we confirmed by immunohistochemistry **(IHC)** against PV at P21, the developmental stage when PV reaches uniform expression in L2/3 ACC (Supplementary Fig. 2). Approximately 83% of tdTomato-positive (**tdTom^+^**) neurons in L2/3 ACC were co-labeled with PV immunolabelling at P21, increasing to 94% in adulthood (Fig. 2B–2D). We also confirmed minimal overlap between S5E2-Tom^+^ cells and Somatostatin (**SST**) expressing interneurons at P15, which are another major source of inhibition to L2/3 PYR (Supplementary Fig. 3). These results confirm the high specificity of S5E2-based reporters for PVIN^62^, and S5E2^+^ neurons will henceforth be referred to as PVIN in this study. We observed no difference in mEPSC amplitude or frequency in ACC L2/3 PVIN at P15 (Fig. 2G and 2H). There was also no difference in mEPSC rise and decay times or membrane capacitance (Supplementary Fig. 4A–4C and 4E), suggesting that loss of Shank3 does not impair the early maturation of AMPAR-mediated current onto ACC L2/3 PVIN.

**Figure 2.**
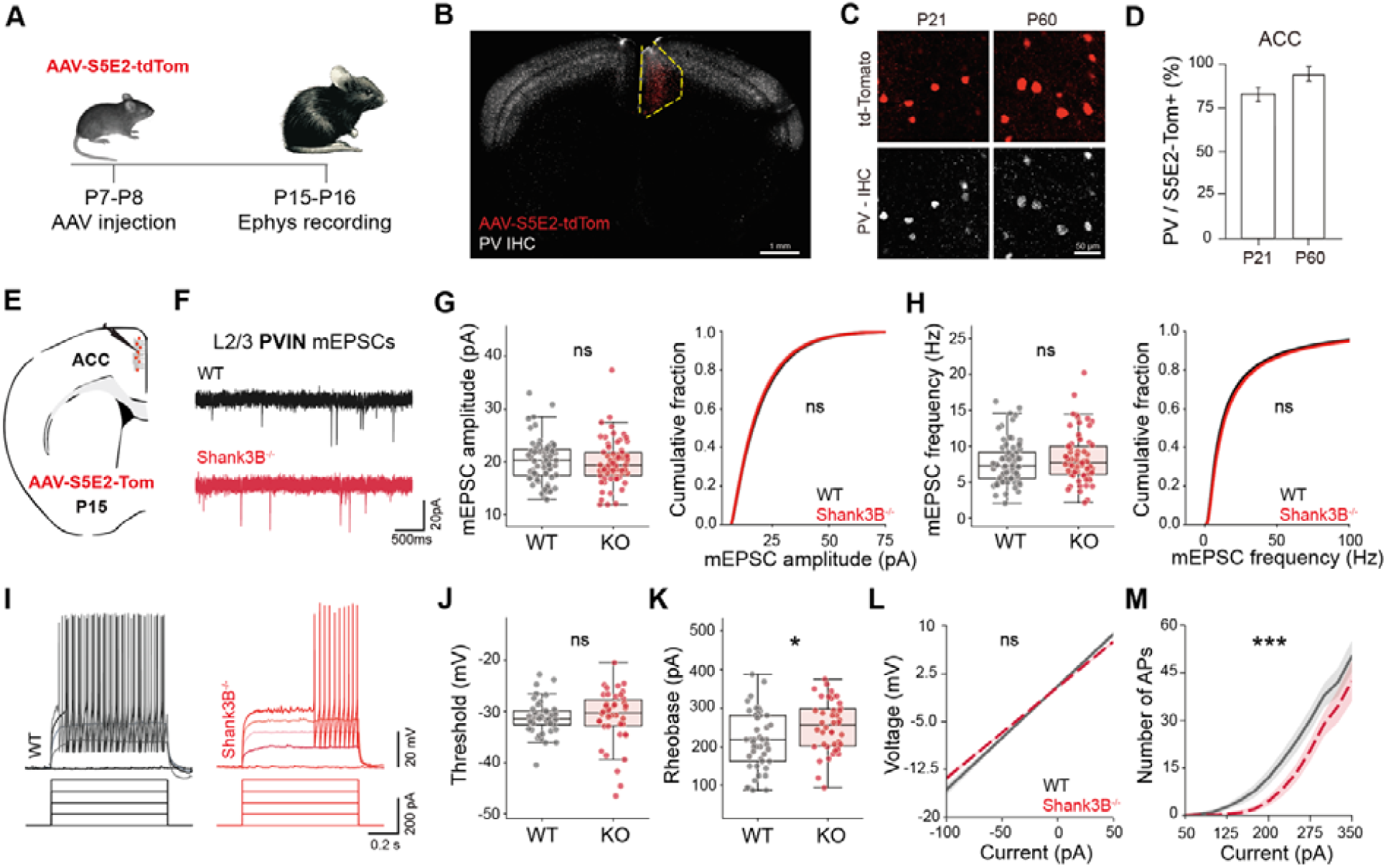
Reduced excitability but normal AMPAR-mediated synaptic inputs of ACC L2/3 PVIN in P15 Shank3B^-/-^ mice. **(A)** Schematic of experimental timeline showing AAV-S5E2-tdTomato injection in WT and Shank3B^-/-^ mice at P7-P8, followed by electrophysiological recordings at P15-P16. **(B)** Representative coronal section showing S5E2-Tom expression and PV immunohistochemical labeling in ACC. Scale bar = 1mm. **(C)** Confocal images of S5E2-tdTom-positive and PV-positive cells in the ACC at P21 and P60, showing high degree of colocalization. Scale bar = 50µm. **(D)** Quantification of the percentage of PV-positive neurons expressing AAV-S5E2-tdTomato in the ACC at P21 and P60 (Average±SEM: P21 82.5±1.9%; P60 94.3±4.1%). **(E)** Schematic representing whole-cell recordings in S5E2^+^ interneurons in L2/3 ACC of P15 mice **(F)** Representative AMPAR mEPSCs in L2/3 ACC S5E2^+^ cells of WT (black) and Shank3B^-/-^ (red) mice. **(G)** Average (left) and cumulative distribution (right) of mEPSC amplitude (Average±SD, WT 20.1±4.1pA; KO 19.8±4.3pA, MW test p=0.54. KS test p=1.0) and **(H)** frequency per neuron (Average±SD, WT 7.8±3.0Hz; KO 8.4±3.5Hz, MW test p=0.35. KS test p=0.98). **(I)** Representative traces of action potentials recorded in response to step current injections in L2/3 ACC S5E2^+^ interneurons of WT (black) and Shank3B^-/-^ (red) mice. **(J)** Average AP threshold voltage (Average±SD, WT -31.2±3.3mV; KO -30.7±5.3mV, MW test p=0.11) and **(K)** Rheobase current (Average±SD, WT 217.1±75.3pA; KO 255.8±65.4pA, MW test p=0.02). **(L)** I-V relationship of L2/3 ACC S5E2^+^ cells (Two-way ANOVA of I-V curve p=0.077). **(M)** I-F relationship of L2/3 ACC S5E2^+^ cells (Two-way ANOVA of I-F curve p=3.3e-8 in genotype). For PV quantification, n=3,3 WT and Shank3B^-/-^ mice at both ages. For mEPSC recording, n=69,65 neurons from n=6,6 WT and Shank3B^-/-^ mice. For current-clamp recording, n=46,39 neurons from n=4,4 WT and Shank3B^-/-^ mice. Box plots show median and quartile values. eCDFs represent average of CDF from mEPSCs of each recorded neuron. Box plots *P<0.05 with MW test. eCDFs *P<0.05 with KS test. Regression plots ***P<0.001 with two-way ANOVA.

To characterize the excitability of PVIN, we performed current-clamp recordings in S5E2^+^ neurons of L2/3 ACC of P15 WT and Shank3B^-/-^ mice. No difference in AP threshold potential and membrane resistance were detected in L2/3 PVIN (Fig. 2J and 2L). However, we observed a significant increase in rheobase current (Fig. 2K) and a rightward shift in the I-F curve, indicating reduced excitability of L2/3 PVIN in Shank3B^-/-^ mice at this early postnatal stage (Fig. 2M). We confirmed that the differences in PVIN intrinsic properties were not driven by an outlier mouse by verifying that no individual mouse average deviated more than 2 standard deviations from the group mean (Supplementary Fig. 5). In addition, despite the reduced rate of AP firing, no changes in RMP or AP kinetics were observed between genotypes (Supplementary Fig. 4F–4L). Together, these findings reveal a selective reduction in L2/3 ACC PVIN excitability in Shank3B^-/-^ mice, specifically affecting rheobase and input–output I-F relationship while sparing most passive membrane properties.

## Loss of Shank3 reduces PVIN-mediated inhibition onto L2/3 PYR at P15

PVIN are critical regulators of cortical circuit activity and provide strong GABAergic inhibition to neighboring PYR^35,36^. To determine whether loss of Shank3 impairs PVIN-dependent inhibition in ACC L2/3 circuits during postnatal development, we compared optogenetically evoked PVIN inhibitory postsynaptic currents (**oIPSCs**) in L2/3 PYR of P15 WT and Shank3B^-/-^ mice. Mice were injected at P7 with AAV9-S5E2-C1V1-YFP in ACC to express the red-shifted excitatory opsin C1V1 in PVIN (Fig. 3A) and acute brain slices containing ACC were used for whole cell-recordings at P15. GABA-mediated currents were isolated by performing voltage-clamp recordings at V_m_= 0mV in the presence of RS-CPP and NBQX, NMDAR and AMPAR blockers, respectively. oIPSCs were evoked using 1ms 563 nm light pulse to stimulate C1V1 expressing fibers. Notably, the amplitude of oIPSC in Shank3B^-/-^ PYR was largely reduced compared to WT, indicating that loss of Shank3 impairs PVIN-mediated inhibition (Fig. 3C). However, no difference in latency to oIPSC peak was detected suggesting normal postsynaptic GABAergic signaling (Fig. 3D). To rule out potential confounds caused by variable C1V1 expression levels or viral targeting efficiency, we regressed the YFP intensity in the field of view of the recorded neuron against its oIPSC amplitude. This analysis revealed consistently lower currents in Shank3B^-/-^ pyramidal neurons compared to WT across all fluorescence intensity levels (Supplementary Fig. 6A), indicating that reduced PVIN-dependent oIPSC amplitude in P15 Shank3B^-/-^ ACC is not caused by reduced opsin expression. To further exclude potential genotype-dependent differences in opsin conductance or membrane trafficking, we assessed AP success rates in PVINs of expressing S5E2-C1V1 using current-clamp recordings, applying the same light stimulation parameters used in oIPSC recordings. We confirmed that these stimuli reliably evoked APs with a 100% success rate across all trials and recorded neurons of both WT and Shank3B^-/-^ mice, indicating no genotype-dependent differences in light-evoked firing in C1V1-expressing PVINs (Supplementary Fig. 6D, 6E, and 6G). Additionally, we measured direct opsin-mediated currents using voltage-clamp recordings in C1V1-expressing cells and similarly found no significant differences between genotypes (Supplementary Fig. 6F), These results indicate that the reduced oIPSC amplitudes recorded in PYR neurons of Shank3B^-/-^ mice are not due to abnormal opsin function in PVIN.

**Figure 3.**
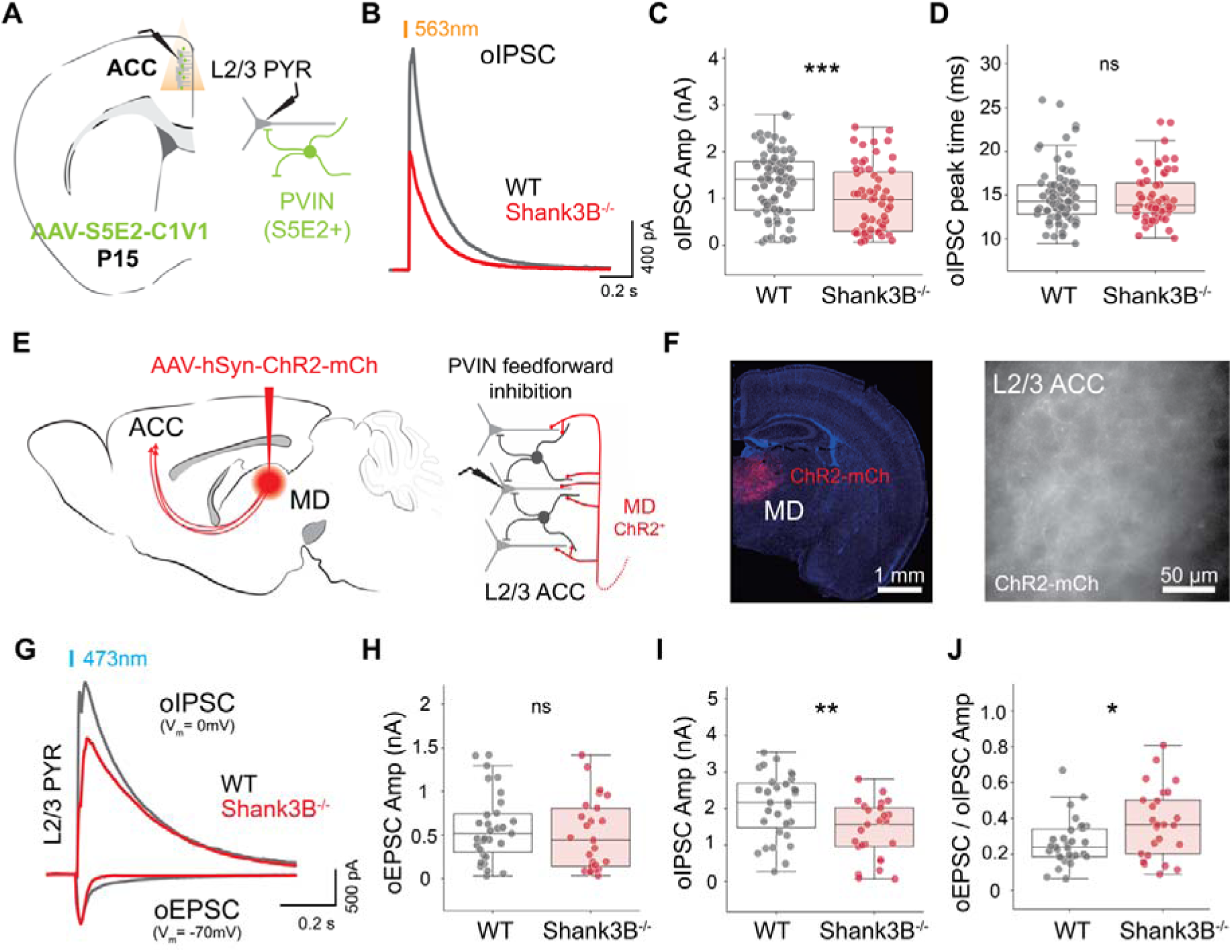
Loss of Shank3 reduces PVIN-mediated inhibition onto ACC L2/3 PYR at P15. **(A)** Schematic representing whole-cell recordings in L2/3 ACC of P15 mice previously injected with AAV-S5E2-C1V1-YFP to express ChR2 in S5E2^+^ interneurons. **(B)** Representative traces of oIPSCs evoked in L2/3 ACC PYR of WT (black) and Shank3B^-/-^ (red) mice in response to optogenetic stimulation of S5E2^+^ interneurons **(C)** Neuron average oIPSC amplitude (Average±SD, WT 1.31±0.67nA; KO 0.98±0.71nA, MW test p=0.004) and **(D)** peak time (Average±SD, WT 14.9±3.3ms; KO 14.8±2.9ms, MW test p=0.95). **(E)** Schematic of the viral targeting strategy for optogenetic activation of projections from the mediodorsal thalamus (MD) to the ACC using AAV8-hSyn-ChR2-mCherry. MD fiber stimulation drives PVIN dependent FFI in L2/3 ACC. **(F)** Representative images of ChR2-mCh expression in P15 MD (left) and L2/3 ACC (right). **(G)** Representative oEPSCs (AMPAR currents) and oIPSCs (GABAR currents) in L2/3 ACC PYR of WT (black) and Shank3B^-/-^ (red) mice. Scale bars = 1mm, left panel; 50µm, left panel. **(H)** Average oEPSC (Average ±SD: WT 581.8±391.8 pA; KO 505.8±409.3 pA, MW test p=0.4) and **(I)** oIPSC peak amplitude (Average±SD, WT 2.07±0.88 nA; KO 1.39±0.78 nA, MW test p=0.007) per neuron. **(J)** Ratio of oEPSC/oIPSC amplitudes calculated per neuron (Average±SD, WT 0.3±0.1; KO 0.4±0.2, MW test p=0.03). For oIPSC recording, n=77,59 neurons from n=7,7 WT and Shank3B^-/-^ mice. For thalamic input recording, n=32,35 neurons from n=4,4 WT and Shank3B^-/-^ mice. Box plots show median and quartile values. Box plots *P<0.05, **P<0.01, ***P<0.001 with MW U test.

PYR and PVIN in L2/3 of the ACC receive robust glutamatergic innervation from the mediodorsal thalamus (**MD**), with MD-derived inputs selectively driving PVIN-dependent feedforward inhibition (**FFI**) in this region^63^. We recorded MD-driven optogenetically induced excitatory postsynaptic currents (**oEPSCs**) and FFI onto L2/3 PYR in ACC of P15 WT and Shank3B^-/-^ mice. To express ChR2 in MD fibers mice were injected AAV8-hSyn-ChR2-mCherry in MD at P7 (Fig. 3E and 3F). Voltage-clamp recordings in L2/3 PYR were performed in ACSF with RS-CPP, using a low chloride Cs^+^ based internal solution with E_Cl_= -70mV. AMPAR and GABA_a_R mediated currents were isolated by holding the recorded neuron at V_m_= -70mV and 0mV, respectively. Interestingly, we observed a significant reduction in MD-driven GABA_a_R oIPSC amplitude, while AMPAR oEPSC amplitude remained unchanged (Fig. 3H and 3I), resulting in an elevated ratio of excitation/inhibition (**E/I**) in Shank3B^-/-^ L2/3 PYR when compared to WT (Fig. 3J). PVIN firing is regulated in part by NMDAR-mediated currents^37,64,65^ and previous studies reported reduced NMDA/AMPA (**N/A**) ratio in cortical neurons of adult Shank3B^-/-^ mice^66^. To determine whether impaired MD-driven FFI in ACC of Shank3B^-/-^ mice is caused by abnormal NMDAR currents onto PVIN, we assessed the N/A ratio of MD-driven oEPSCs recorded at -70mV and +40mV onto L2/3 PVIN and PYR. We observed no significant differences between genotypes for both cell types (Supplementary Fig. 7), indicating normal N/A ratio of MD thalamocortical synapses in L2/3 ACC of Shank3B^-/-^ mice at this developmental stage, and suggesting that reduced FFI is unlikely to result from altered NMDAR-mediated currents in PVIN. Taken together, these results indicate that PVIN-dependent inhibition onto L2/3 PYR in the ACC is markedly reduced by the loss of Shank3 as early as P15.

## Reduced HCN currents in ACC L2/3 PVIN and PYR of P15 Shank3B^-/-^ mice

The rightward shift in the I–F relationship in Shank3B^-/-^ ACC PVINs (Fig. 2M), despite normal synaptic input and passive membrane properties, suggests a disruption in active mechanisms regulating intrinsic excitability and action potential firing. Previous studies have shown that loss of Shank3 in mice leads to HCN channelopathy in thalamic neurons^46,48^. Given the critical role of HCN channels in regulating burst firing dynamics, we assessed whether HCN-mediated I_h_ currents are already impaired in ACC L2/3 PVINs and PYRs of Shank3B^-/-^ mice at P15. To isolate HCN mediated currents we performed whole-cell recordings in voltage-clamp with potassium based internal solution including K^+^ channel blocker tetraethylammonium (**TEA**) and extracellular bath application of 4-AP, TTX and BaCl_2_ and measured current changes in response to hyperpolarizing voltage steps (Fig. 4A). Incubation with HCN channel blocker ZD7288 abrogated all currents recorded in PVINs under these conditions, confirming the presence of soma-recorded I_h_ currents in L2/3 ACC PVINs at this age, consistent with previous developmental studies in basket cells of somatosensory cortex in mice^67^ (Fig. 4B). We found a significant reduction in I_h_ current density in both L2/3 PVINs and PYRs of Shank3B^-/-^ mice compared to WT controls (Fig. 4C–E, 4G–I). Sigmoidal fits of the voltage–current density relationship revealed a reduction in maximum I_h_ current density without changes in slope, suggesting a decrease in HCN channel number, but normal voltage-dependent gating. In addition, we observed a non-statistically significant trend toward faster decay tau of PVIN I_h_ currents in Shank3B^-/-^ mice, whereas I_h_ decay kinetics in PYRs were similar between genotypes (Fig. 4F and 4J). Together, these findings indicate that loss of Shank3 reduces HCN channel currents in both PVINs and PYRs in L2/3 ACC as early as P15, supporting the idea that HCN channelopathy in Shank3B^-/-^ mice emerges during early development.

**Figure 4.**
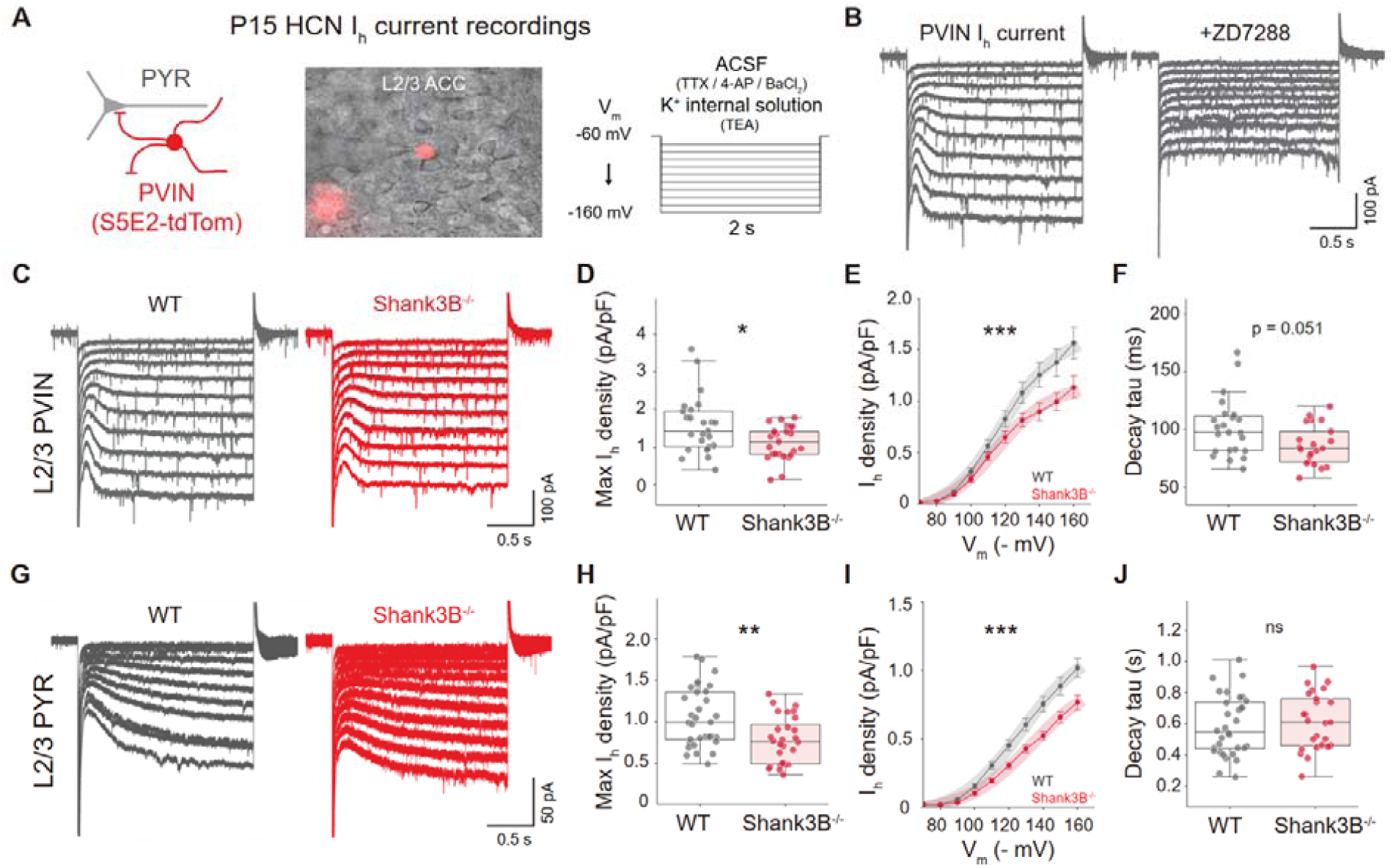
Reduced hyperpolarization-activated cation currents of ACC L2/3 PVIN and PYR in P15 Shank3B^-/-^ mice. **(A)** Schematic condition for Ih current recordings in L2/3 ACC S5E2^+^ interneurons and PYR by performing voltage-clamp recording with hyperpolarizing voltage steps. **(B)** Representative traces of I_h_ current recordings in response to hyperpolarizing voltage steps in WT L2/3 ACC S5E2^+^ interneurons by recording with the bath and internal solution adding without or with ZD7288. **(C)** Representative traces of I_h_ current recordings in response to hyperpolarizing voltage steps in L2/3 ACC S5E2^+^ interneurons of WT (black) and Shank3B^-/-^ (red). **(D)** Maximum I_h_ current density (Average±SD, WT 1.6±0.8pA; KO 1.1±0.4pA, MW test p=0.03) and **(E)** Current density/voltage relationship of Ih currents in L2/3 ACC S5E2^+^ interneurons (Two-way ANOVA, p=7.4e-8 in genotype, sigmoid curve fit, WT Max density:1.741, slope:0.062; KO Max current density:1.231, slope:0.064). **(F)** I_h_ current decay tau in L2/3 ACC S5E2^+^ interneurons (Average±SD, WT 102.1±25.4ms; KO 86.8±17.1ms, MW test p=0.051). **(G)** Representative traces of I_h_ current recordings in response to hyperpolarizing voltage steps in L2/3 ACC PYR of WT (black) and Shank3B^-/-^ (red). **(H)** Maximum Ih current density (Average±SD, WT 1.1±0.4pA; KO 0.8±0.3pA, MW test p=0.003) and **(I)** Current density/voltage relationship of I_h_ currents in L2/3 ACC PYR (Two-way ANOVA, p=2.2e-13 in genotype, sigmoid curve fit, WT Max density:1.253, slope:0.054; KO Max density:0.987, slope:0.053). **(K)** Ih current decay tau in L2/3 ACC PYR (Average±SD, WT 585.0±193.0ms; KO 614.6±183.9ms, MW test p=0.484). n=25,25 S5E2^+^ interneurons from n=4,4 WT and Shank3B^-/-^ mice; n=32,28 PYR from n=6,5 WT and Shank3B^-/-^ mice. Box plots show median and quartile values. Box plots *P<0.05 **P<0.01 with MW test. Current density/voltage relationship are shown as mean, sem and sigmoid curve fit (translucent thick line), ***P<0.001 with two-way ANOVA.

## Reduced AMPAR-mediated synaptic input and excitability of ACC L2/3 PYR in adult Shank3B^-^**^/-^ mice.**

To investigate late-stage adaptations in excitatory synaptic function and neuronal excitability, we characterized AMPAR-mediated mEPSCs and the excitability of L2/3 PYR neurons in adult Shank3B^-/-^ mice (Fig. 5A-B). ACC L2/3 PYR displayed lower mEPSC amplitude, consistent to previous reports^53^ (Fig. 5C), but no difference in mEPSC frequency was observed (Fig. 5D). In addition, adult Shank3B^-/-^ mice displayed longer mEPSC rise time and slower mEPSC rise velocity of L2/3 PYR in ACC (Fig. 5E and 5F), suggesting slower neurotransmission potentially due to abnormal synaptic structural coupling or AMPAR localization. mEPSC decay tau and membrane capacitance were, however, unaffected, suggesting these phenotypes are not due to major changes in AMPAR conductance or cell size (Supplementary Fig. 8A–8C).

**Figure 5.**
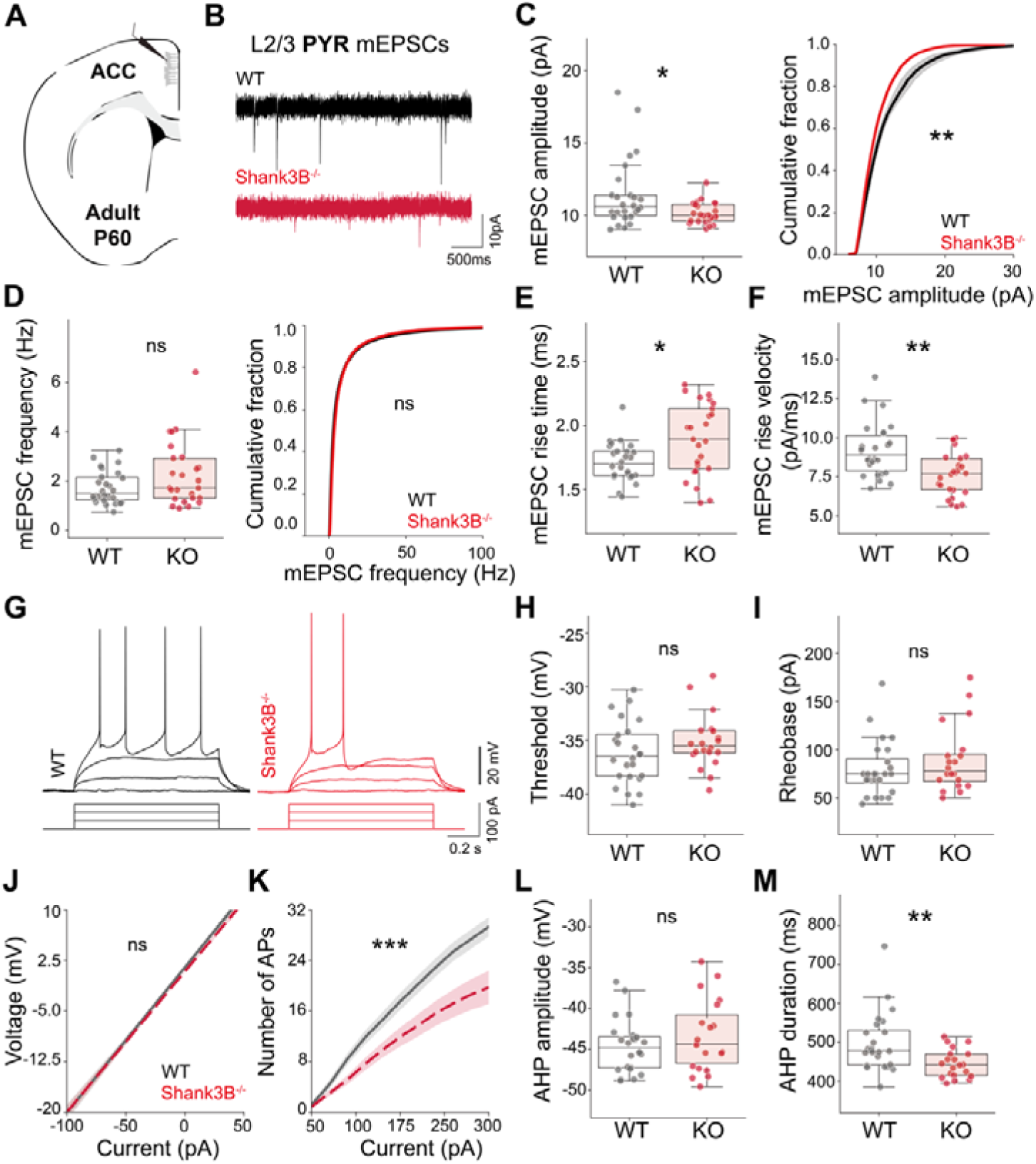
Reduced AMPAR-mediated synaptic strength and hypoexcitability of ACC L2/3 PYR in Adult Shank3B^-/-^ mice. **(A)** Schematic representing coronal brain section and whole-cell recordings of L2/3 PYR in ACC of P60 mice. **(B)** Representative AMPAR mEPSCs recorded in L2/3 ACC PYR of WT (black) and Shank3B^-/-^ (red) mice. **(C)** Average (left) and cumulative distribution (right) of mEPSC amplitude (Average±SD, WT 11.4±2.4pA; KO 10.1±1.7pA, MW test p=0.04. KS test p=0.002) and **(D)** frequency per neuron (Average ±SD, WT 1.7±0.7 Hz; KO 2.3±1.3 Hz, MW test p=0.22. KS test p=0.07). **(E)** Average mEPSC rise time (Average±SD, WT 1.7±0.2ms; KO 1.9±0.3 ms, MW test p=0.01) and **(F)** velocity per neuron (Average ±SD, WT 9.2±1.7 pA/ms; KO 7.7±1.4 pA/ms, MW test p=0.003) showing slower mEPSC kinetics in adult L2/3 ACC PYR of Shank3B^-/-^ mice. **(G)** Representative current-clamp recordings in L2/3 ACC PYR of WT (black) and Shank3B^-/-^ (red) mice. **(H)** Average of AP threshold (Average±SD, WT -36.2±2.9 mV; KO -35.1±2.5 mV, MW test p=0.18). **(I)** Rheobase current per neuron (Average±SD, WT 79.9±29.2 pA; KO 89.7±34.6 pA, MW test p=0.35). **(J)** I-V (Two-way ANOVA of I-V curve p=0.22 in genotype) and **(K)** I-F relationship of L2/3 ACC PYR (Two-way ANOVA of I-F curve p=3.8e-17 in genotype). **(L)** Average afterhyperpolarization current amplitude (Average±SD, WT -44.6±3.3 mV; KO -43.5±4.4 mV, MW test p=0.49) and **(M)** duration per neuron (Average ±SD, WT 498.9±76.1 ms; KO 444.9±36.6 ms, MW test p=0.006). For mEPSC recording, n=26,25 neurons from n=3,3 WT and Shank3B^-/-^ mice. For current-clamp recording, n=24,20 neurons from n=3,3 WT and Shank3B^-/-^ mice. Box plots show median and quartile values. eCDFs represent average of CDF from mEPSCs of each recorded neuron. Box plots *P<0.05, **P<0.01 with MW test. eCDFs **P<0.01 with KS test. Regression plots ***P<0.001 with two-way ANOVA.

Next, we examined the intrinsic properties of L2/3 PYR in adult ACC by current-clamp recordings. No differences were observed between genotypes in AP threshold, rheobase, or membrane resistance (Fig. 5G–5J). However, Shank3B^-/-^ PYR exhibited a significant reduction in the I–F relationship, indicating reduced excitability (Fig. 5K). Conversely, no difference in RMP and AP kinetics were found, suggesting normal AP generation (Supplementary Fig. 8D and 8E–8H), but the duration of the AHP was significantly reduced (Fig. 5L and 5M). Together, these results suggest that loss of Shank3 leads to long-term adaptations in frontal cortical circuits, marked by reduced AMPAR-mediated synaptic strength and decreased excitability of ACC L2/3 PYRs in adulthood.

## Late-stage adaptations in AMPAR-mediated synaptic input and intrinsic excitability in L2/3 ACC PVIN of adult Shank3B^-/-^ mice

GABAergic interneurons of the primary somatosensory cortex of adult Shank3B^-/-^ mice exhibit dampened responses to peripheral stimulation^54^, but whether PVIN are specifically affected remains unknown. Given the excitability deficits observed in ACC L2/3 PVIN at P15 (Fig. 2), we assessed whether the loss of Shank3 leads to permanent PVIN functional deficits in adulthood. To address this, we first characterized AMPAR-dependent mEPSCs in ACC L2/3 PVIN of mice injected with AAV5-S5E2-tdTom two weeks prior to recordings (Fig. 6A and 6B). We observed a significant reduction of both mEPSC amplitude and frequency in PVIN of adult Shank3B^-/-^ mice when compared to WT controls (Fig. 6C and 6D). No differences were observed in mEPSC rise and decay times, or membrane capacitance, but mEPSC rise velocity was increased (Supplementary Fig. 9A–9C and 9E). Together, these findings suggest reduced number and strength of AMPAR-mediated synaptic inputs onto ACC L2/3 PVIN in adult Shank3B^-/-^ mice.

**Figure 6.**
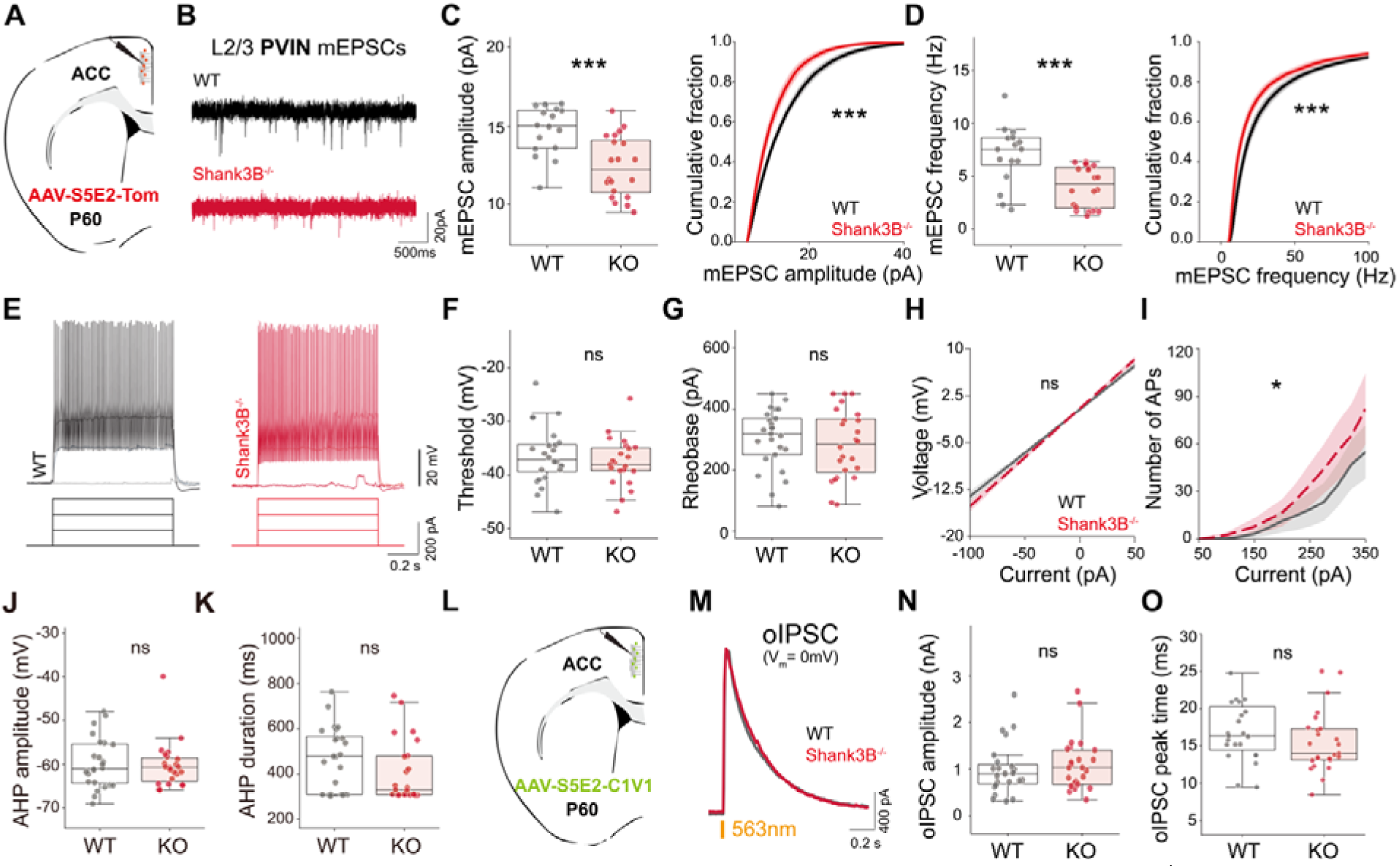
Adult-stage adaptations in synaptic and intrinsic excitability in L2/3 ACC PVIN of adult Shank3B^-/-^ mice. **(A)** Schematic representing whole-cell recordings in S5E2^+^ interneurons in L2/3 ACC of P60 mice **(B)** Representative AMPAR mEPSCs in L2/3 ACC S5E2^+^ cells of WT (black) and Shank3B^-/-^ (red) mice. **(C)** Average (left) and cumulative distribution (right) of mEPSC amplitude (Average±SD, WT 14.9±1.6 pA; KO 12.5±2.0 pA, MW test p=9.8e-4. KS test p=8e-5) and **(D)** frequency per neuron (Average±SD, WT 7.1±2.6 Hz; KO 4.4±1.8 Hz, MW test p=5e-4. KS test p=1e-5). **(E)** Representative traces of current-clamp recordings in L2/3 ACC S5E2^+^ interneurons of WT (black) and Shank3B^-/-^ (red) mice. **(F)** Average of AP threshold potential (Average±SD, WT -36.4± 5.4mV; KO -37.7±4.5 mV, MW test p=0.46) **(G)** and rheobase current (Average±SD, WT 301.9±97.1 pA; KO 283.6±112.3 pA, MW test p=0.56). **(H)** I-V (Two-way ANOVA of I-V curve p=0.28 in genotype) and **(I)** I-F relationship of L2/3 ACC S5E2^+^ cells (Two-way ANOVA of I-F curve p=0.015 in genotype). **(J)** Average afterhyperpolarization current amplitude (Average±SD, WT 59.8±5.8 mV; KO 60.2±5.4 mV, MW test p=0.89) and **(K)** duration per neuron (Average±SD, WT 466.8±141.6 ms; KO 408.1±139.2 ms, MW test p=0.18). **(L)** Schematic representing coronal brain section and whole-cell recordings in L2/3 ACC on adult mice previously injected with AAV-S5E2-C1V1-YFP for optogenetic stimulation of S5E2^+^ interneurons. **(M)** Representative S5E2-dependent oIPSCs in L2/3 ACC PYR of adult WT (black) and Shank3B^-/-^ (WT) mice. **(N)** Average oIPSC amplitude and (Average±SD, WT 1.00±0.56 nA; KO 1.09±0.56 nA, MW test p=0.54. **(O)** and oIPSC peak time (Average±SD, WT 17.7±6.4 ms; KO 15.5±4.1 ms, MW test p=0.14). For S5E2^+^ cells mEPSC recordings, n=30,29 neurons from n=4,4 WT and Shank3B^-/-^ mice. For S5E2^+^ cells current-clamp recordings, n=25,24 neurons from n=4,4 WT and Shank3B^-/-^ mice. For oIPSC recordings, n=23,25 neurons from n=4,4 WT and Shank3B^-/-^ mice. Box plots show median and quartile values. eCDFs represent average of CDF from mEPSCs of each recorded neuron. Box plots ***P<0.001 with MW test. eCDFs ***P<0.0001 with KS test. Regression plots *P<0.05 with two-way ANOVA.

To determine whether the deficits in neuronal excitability observed at P15 (Fig. 2) are maintained in adulthood, we characterized PVIN intrinsic excitability at P60. Surprisingly, and contrary to what we observed during postnatal development, no change in AP threshold potential, rheobase current, or membrane resistance were detected in ACC L2/3 PVIN in adult Shank3B^-/-^ mice (Fig. 6E–6H). No change of resting membrane potential and AP properties were also found (Supplementary Fig. 9F– 9J). However, PVIN displayed increased AP firing in response to current injections, which is opposite to the phenotype observed at P15 (Fig. 6I). Likewise, ACC L2/3 PYR of Shank3B^-/-^ mice showed no differences in PVIN-dependent oIPSC amplitude or peak time (Fig. 6L–6O and Supplementary Fig. 6B). These results reveal complex developmental trajectories of PVIN synaptic and intrinsic properties and suggest extensive compensatory mechanisms in PVIN function throughout development.

Decreased PV expression has been reported in various brain areas in adult Shank3B^-/-^ mice^26,27,32^. To examine whether PV expression is altered in ACC and anterior (upper lip/ forelimb) somatosensory cortex (**S1**) of adult Shank3B^-/-^ mice, we analyzed the density of PVIN and PV expression by IHC in L2/3 and L5 ACC and S1 in P60 mice. Interestingly, we observed no differences in PV^+^ neuron density or PV fluorescence signal intensity between genotypes in any of the cortical layers of ACC or S1 (Supplementary Fig. 10), suggesting normal PV expression and PVIN cell number in frontal cortical regions of Shank3B^-/-^ mice.

## Reduced frequency of Ca^2+^ transients in ACC PVIN of P15-P17 Shank3B^-/-^ mice

To test whether the excitability and connectivity deficits observed in ACC PVIN at P15 (Fig. 2 and 3) are related with abnormal patterns of cortical activity, we measured spontaneous neuronal activity in dorsal ACC of awake WT and Shank3B^-/-^ mice at P15-P17 by *in vivo* two-photon calcium imaging. Mice were co-injected at P7 with AAV9-Syn-GCaMP8s and AAV5-S5E2-tdTom to express GCaMP8s in all cortical neurons and tdTomato in PVIN for labeling (Fig. 7A and 7B). Mice were implanted with a cranial window at P15-P17 and imaged in the same day. Spike-related calcium **(Ca^2+^)** fluorescence of GCaMP8s expressing neurons was recorded at 15Hz frame rate with 920nm excitation light and analyzed using Suite2P^68^. Regions of interest **(ROI)** from PVIN were identified based on tdTomato expression using CellPose^69^. GCaMP8s activity traces from the majority of putative PYR (S5E2-tdTom negative) was characterized by discrete and sparse Ca^2+^ transients, while those from putative PVIN (S5E2-Tom positive) exhibited high frequency, low amplitude transients (Fig. 7C), consistent with the high firing rates and burst firing of cortical PVIN *in vivo*^64^. To compare calcium transient properties across animals we calculated ΔF/F activity and analyzed individual Ca^2+^ events per ROI (see Methods). We detected no significant genotype differences in the average frequency or amplitude of Ca^2+^ events in PYR, suggesting normal activity patterns of ACC L2/3 PYR at P16 (Fig. 7D and 7E). In contrast, changes in PVIN Ca²⁺ activity were more pronounced, showing a significant reduction in event frequency but no difference in event amplitude (Fig. 7G and 7H). We also calculated the area under the curve **(AUC)** of individual Ca^2+^ events and found no difference in AUC of both putative PYR and PVIN in ACC of P16 Shank3B^-/-^ mice (Fig. 7F and 7I). Although the average PYR Ca^2+^ event frequency did not show significant differences between genotypes, empirical cumulative distribution function **(eCDF)** analysis of event frequency distribution indicated a trend toward hypoactivity in PYR (Fig. 7D). However, further analysis using deconvolved ΔF/F data to assess event frequency did not support this finding and showed no significant differences in either the mouse averages or the eCDF of deconvolved impulse frequency in PYR (Supplementary Fig. 11A). In contrast, analysis of deconvolved ΔF/F activity traces of PVIN confirmed a significant reduction in frequency of Ca²⁺ transients, consistent with lower ACC PVIN activity (Supplementary Fig. 11E). No differences in Ca^2+^ transient rise time, duration, and decay tau were detected in both PYR and PVIN between genotypes (Supplementary Fig. 11B–11D and 11F– 11H). Together, these results suggest that loss of Shank3 causes reduced activity of PVIN in ACC during early postnatal development.

**Figure 7.**
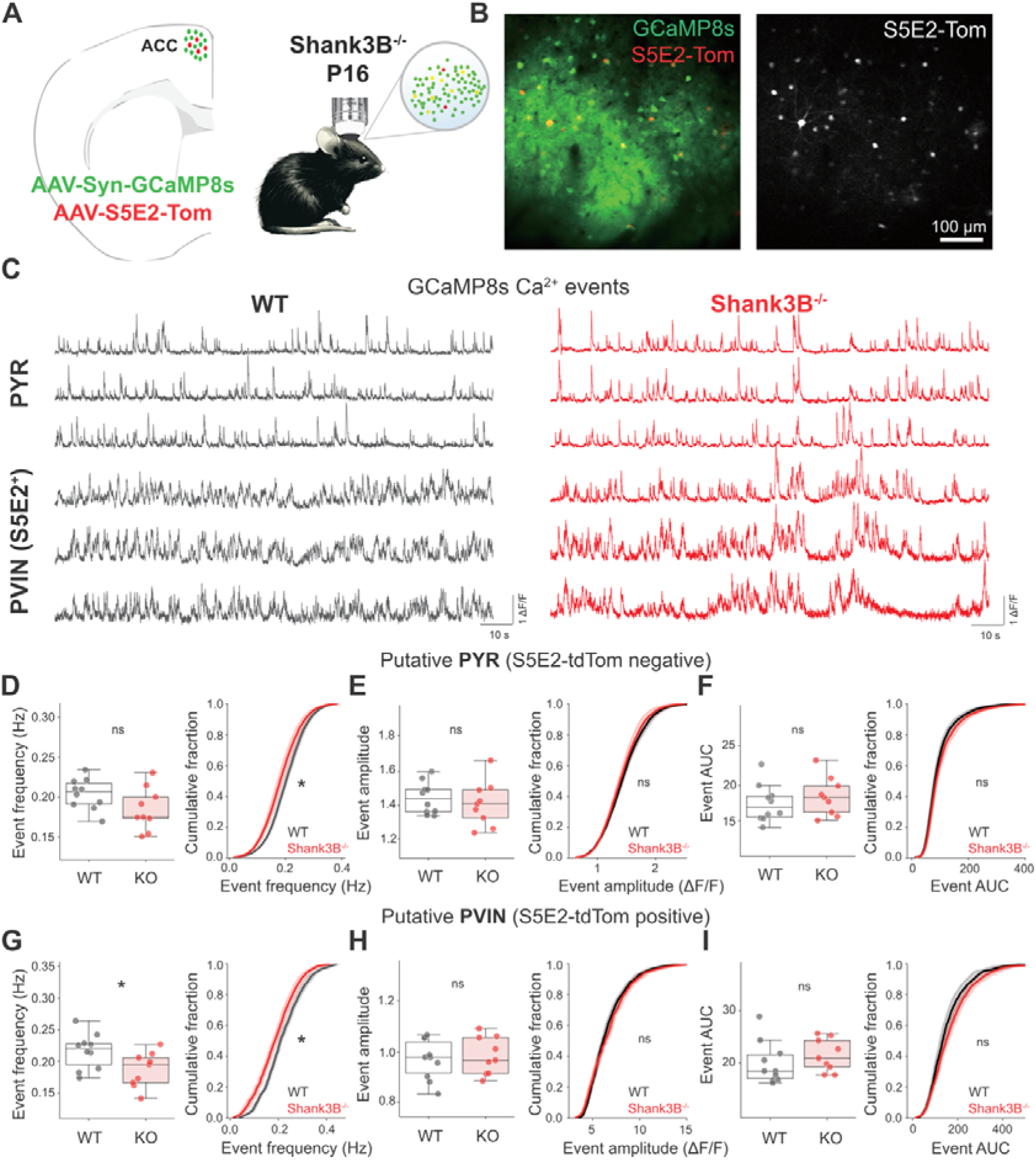
Reduced frequency of Ca^2+^ transients in ACC PVIN of P15-P17 Shank3B^-/-^ mice. **(A)** Schematic diagram of in vivo calcium imaging in dorsal ACC of mouse co-injected at P7 with AAV9-hSyn-GCaMP8s and AAV5-S5E2-tdTom in dACC and imaged at P15-P17 in a 2-photon microscope through a cranial window. **(B)** Fluorescent images showing co-expression of GCaMP8s (green) and S5E2-tdTomato (red) in the dACC. Right panel shows S5E2-tdTomato positive interneurons. Scale bar = 100 µm. **(C)** Representative traces of GCaMP8s calcium events from putative pyramidal (S5E2-tdTom negative) neurons and PVIN (S5E2-Tom positive) neurons in WT (black) and KO (red) mice. **(D)** Average (left) and cumulative distribution (right) of GCaMP8s event frequency (Average±SD, WT 0.20±0.2Hz; KO 0.19±0.03Hz, MW test p=0.13, KS test p=0.03) **(E)** Average (left) and cumulative distribution (right) of GCaMP8s event amplitude (Average±SD, WT 1.45±0.09; KO 1.42±0.13, MW test p=0.60, KS test p=0.97, n=10,9 WT and Shank3B^-/-^ mice). **(F)** Event area under the curve of putative PYR (Average±SD, PYR event AUC, WT 17.4±2.4; KO 18.3±2.5, MW test p=0.39, KS test p=0.76) **(G)** Average (left) and cumulative distribution (right) of GCaMP8s event frequency (Average±SD, WT 0.20±0.2Hz; KO 0.19±0.03Hz, MW test p=0.04, KS test p=0.02) **(H)** Average (left) and cumulative distribution (right) of GCaMP8s event amplitude (Average±SD, WT 0.97±0.08; KO 0.99±0.07, MW test p=0.65, KS test p=0.61). **(I)** Event area under the curve of putative PVIN (Average±SD, WT 20.1±4.0; KO 21.6±3.1, MW test p=0.27, KS test p=0.16). n=10,9 WT and Shank3B^-/-^ mice. Box plots show median and quartile values. Box plots *P<0.05 with MW test. eCDFs *P<0.05 with KS test.

Deep layer cortical neurons exhibit one of the highest co-expression rates of ASD risk genes^57^ and are functionally impaired in several mice carrying ASD-associated genetic mutations^12,14,15^. To determine whether the deficits in PVIN Ca^2+^ activity are restricted to specific cortical layers we analyzed ROIs segregated by their mediolateral **(ML)** distance to the central sinus corresponding to superficial (100-375 µm) and deep layers (>375 µm). Interestingly, PVIN showed similar trends in event frequency, amplitude, and AUC across the ML axis, suggesting that loss of Shank3 similarly disrupts PVIN activity across cortical layers in dorsal ACC (Supplementary Fig. 12 and 13). To test whether loss of Shank3 alters activity correlation between different cell populations in ACC at P16, we measured Pearson correlation coefficients **(PCC)** of deconvolved ΔF/F activity between PYR-PYR, PYR-PVIN, and PVIN-PVIN ROI pairs. We detected a significant reduction in correlation between all neuronal types with increased cell-cell distance. However, there was no statistical difference in the correlation of Ca^2+^ activity between any cell types across genotypes (Supplementary Fig 12), suggesting that the correlation of ACC PYR and PVIN based on GCaMP8s imaging is not significantly affected by loss of Shank3.

## Discussion

In this study, we investigated the postnatal maturation of PVIN in L2/3 of the ACC in male Shank3B^-/-^ mice, focusing on their intrinsic excitability, glutamatergic synaptic connectivity, and *in vivo* activity. Our results reveal that deficits in L2/3 PVIN excitability are already observed by P15 (Fig. 2), a critical developmental stage analogous to early childhood in humans, when ASD symptoms typically begin to emerge^70^. PVIN hypoexcitability was accompanied by significant impairments in PVIN-mediated inhibition of PYR in L2/3 ACC (Fig. 3) as well as a reduction in HCN-mediated I_h_ currents (Fig. 4). Furthermore, two-photon calcium imaging revealed overall hypoactivity of ACC PVIN during early postnatal development, indicating that the observed physiological abnormalities are associated with PVIN dysfunction *in vivo* (Fig. 7). At P15, L2/3 ACC PYR exhibited normal excitability and glutamatergic synaptic input but already showed a reduction in I_h_ currents, suggesting that HCN channel dysfunction in one of the earliest physiological abnormalities affecting both PYR and PVIN. By adulthood, both cell types underwent substantial phenotypic changes, exhibiting markedly different synaptic and excitability phenotypes compared to early postnatal stages (Fig. 5 and 6). Together, these results indicate that the loss of Shank3 leads to early circuit dysfunction in the ACC and that reduced PVIN excitability and PVIN-mediated inhibition are relevant phenotypes during postnatal cortical maturation in Shank3B^-/-^ mice (Fig. 8).

**Figure 8.**
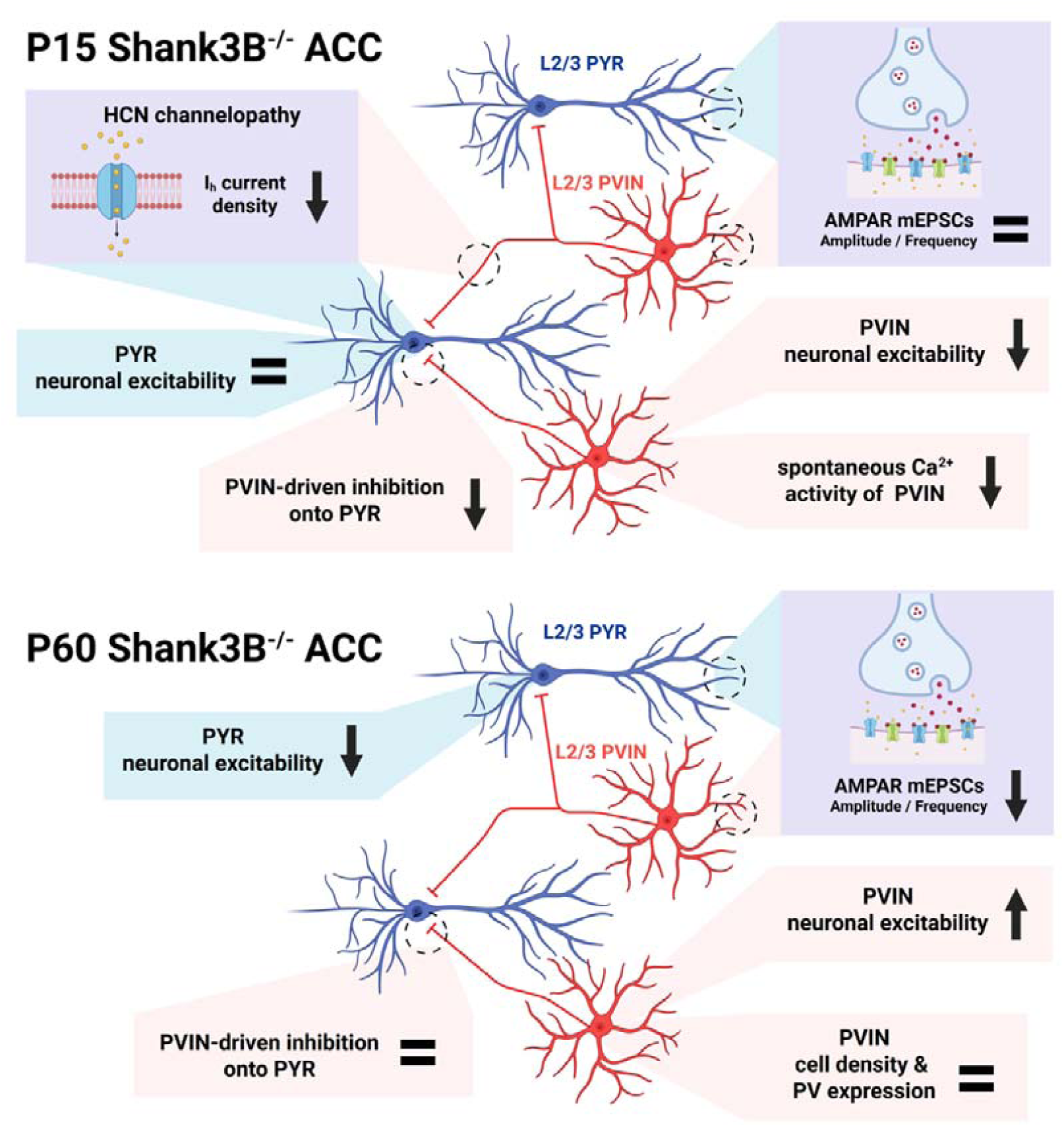
Summary of L2/3 ACC neuron deficits in P15 and P60 Shank3B^-/-^ mice. Both L2/3 PYR (blue) and PVIN (red) show normal AMPAR-mediated mEPSCs in ACC of P15 Shank3B^-/-^ mice. PVIN, but not PYR, show reduced excitability and spontaneous calcium activity *in vivo*. PVIN-dependent inhibition onto PYR is also impaired by loss of Shank3 at this age. Notably, both cell types display significant reductions in HCN-mediated Ih currents, indicating early-onset HCN channelopathy. By adulthood (P60), reduced AMPAR-mediated mEPSCs are observed in both PYR and PVIN. The hypoexcitability of PVIN is reversed, while PYR exhibit hypoexcitability. In contrast, overall current amplitude of PVIN-dependent inhibition onto PYR is restored. The PVIN density and PV expression in L2/3 ACC is also comparable in adult Shank3B^-/-^ mice. This model underscores a critical role for Shank3 in regulating neuronal excitability and HCN channel function during early development, with synaptic dysfunction emerging at later stages. Phenotypes of L2/3 PVIN and PYR are indicated by pink and blue backgrounds, respectively. Shared phenotypes are shown with a purple background.

## Deficits in ACC L2/3 PVIN inhibition emerge during early postnatal development

PVIN hypoexcitability was characterized by elevated rheobase and reduced AP firing in response to depolarizing current injections (Fig. 2). However, other intrinsic properties such as resting membrane potential, membrane resistance, AP threshold, and AP kinetics, were normal (Supplementary Fig. 4), suggesting that the observed PVIN excitability deficits arise from specific impairments in mechanisms related with AP firing^71^. This is consistent with reduced PVIN burst firing observed in Shank2-deficient mice^64^ and support the notion that disruptions to PVIN excitability may represent a shared phenotype across Shank related neurodevelopmental disorders. PVIN-mediated inhibition onto PYR was significantly reduced, both following direct optogenetic stimulation of PVIN and in response to thalamocortical afferent input from MD (Fig. 3). Notably, FFI elicited by MD projections in the ACC is exclusively mediated by PVIN^63^ and is essential for maintaining the balance between excitation and inhibition of cortical circuits^35,36^. These deficits in inhibition are unlikely to be driven by reduced glutamatergic input to PVINs, as both mEPSC properties (Fig. 2E– 2H) and the AMPA/NMDA ratio of MD afferent synapses (Supplementary Fig. 7) were unchanged. PVIN density in both layer 2/3 and layer 5 of the ACC was also comparable between genotypes (Supplementary Fig. 10), strongly suggesting that the deficits in PVIN-mediated inhibition observed at P15 are not caused by a reduction in PVIN cell number. Furthermore, to control for potential artifacts related to variable opsin activation that could confound oIPSC measurements (Fig. 3A–3D), we examined direct opsin-evoked currents and action potential firing in PVINs and found no differences between genotypes (Supplementary Fig. 6D–6G). Together, these results demonstrate that Shank3 deletion already impairs PVIN-mediated inhibition in the L2/3 ACC during early postnatal development and suggest that this dysfunction arises from specific deficits in PVIN membrane excitability and action potential firing, rather than alterations in excitatory input or cell number.

## Early HCN channelopathy in PVIN and PYR of L2/3 ACC in Shank3B^-/-^ mice

HCN mediated I_h_ current density was significantly reduced in both PVIN and PYR of L2/3 ACC in Shank3B^-/-^ mice at P15 (Fig. 4). Notably, reduced I_h_ currents have previously been reported in developing thalamocortical neurons of adult Shank3 knockout **(KO)** mice^46,48,71^, and in human embryonic stem cell-derived neurons with SHANK3 deletions^45^. Together, these findings suggest that Shank3 loss disrupts HCN channel function across multiple neuronal populations, with deficits emerging during early postnatal development. Although we did not directly test whether reduced I_h_ underlies the observed PVIN hypoexcitability, HCN dysfunction provides a plausible mechanistic link between Shank3 loss and impaired PVIN inhibition. In hippocampal basket cells, HCN2 channels regulate AP generation and burst firing^71^. This function aligns with our findings of reduced PVIN-mediated oIPSC amplitude (Fig. 3A–3D) and a rightward-shifted input-output (I-F) relationship (Fig. 2M), despite normal passive membrane properties. HCN channels have also been detected at presynaptic terminals of basket cells in the hippocampus and cerebellum^72–74^ where they facilitate GABA release^75,76^. A similar presynaptic role in cortical PVIN could further contribute to the observed reduction in inhibition, although the expression of presynaptic HCN channels in ACC PVINs remains to be established. Notably, PVIN undergo extensive maturation of intrinsic properties during this postnatal time window^77^. This raises the possibility that reduced I_h_ currents observed in Shank3B^-/-^ PVIN may reflect delayed maturation. However, the preservation of other intrinsic properties, such as resting membrane potential, input resistance, and AP kinetics, argues against a generalized developmental delay. Moreover, many aspects of PVIN maturation, including HCN channel expression and trafficking, are regulated by neural activity^78^, suggesting that the observed excitability deficits might arise as secondary adaptations to altered network activity. However, interneuron-specific deletion of *Shank3* is sufficient to cause cortical circuit dysfunction and behavioral abnormalities in adult mice^54,79,80^, supporting a direct contribution of SHANK3 deficiency within PVINs. Further studies dissecting the cell-autonomous molecular pathways affected by *Shank3* loss in PVINs will be essential for clarifying the mechanisms driving early inhibitory dysfunction and for identifying potential therapeutic targets in SHANK3-associated neurodevelopmental disorders.

## Altered *In Vivo* Calcium Dynamics in ACC L2/3 PVIN

*In vivo* two-photon calcium imaging revealed a significant reduction in the frequency of GCaMP8s calcium transients in PVIN of the ACC in Shank3B^-/-^ mice during early postnatal development (Fig. 7). This reduction in PVIN activity was consistent across the medio-lateral extent of the ACC (Supplementary Fig. 12), indicating widespread hypoactivity across superficial and deep cortical layers. While empirical cumulative distribution function (eCDF) analysis indicated a trend toward reduced event frequency in PYRs, this was not supported by either mouse-level averages or deconvolved impulse frequency analysis of ΔF/F traces. In contrast, deconvolved data showed a significant reduction in calcium event frequency in PVINs, but not PYRs, confirming cell-type-specific hypoactivity at P15 (Supplementary Fig. 11). No significant changes were detected in calcium activity correlations between neuron types, though a subtle trend toward reduced PVIN– PVIN correlation was observed (Supplementary Fig. 12 and 13), suggesting potential alterations in PVIN coupling or network synchrony. However, higher temporal resolution techniques will be needed to resolve these dynamics more conclusively. The observed reduction in ACC PVIN activity is consistent with previous studies reporting reduced frontal cortical activity in Shank3B^-/-^ mice^53,81^, but contrasts with findings of increased neural activity in sensorimotor regions of Shank3-deficient mice during both early postnatal and adult stages^48,54,82–84^. This regional discrepancy may reflect divergent effects of Shank3 loss on frontal versus neocortical circuits, potentially driven by differential developmental trajectories, or varying degrees of afferent or sensory input disruption, which are known to be altered in Shank3B^-/-^ mice^27,31^. The reduction in GCaMP8s transients in PVINs is also consistent with the excitability deficits and reduced AP firing we observed in our whole-cell recordings (Fig. 2), supporting an overall hypoactivity of PVIN in ACC at P15. We observed no differences in AMPAR mEPSC frequency or amplitude or in the AMPAR/NMDAR ratio of thalamocortical inputs to L2/3 ACC PYR at this age. However, it is important to consider the possibility that the early reduction in PVIN activity might reflect a compensatory response to lower afferent drive to frontal circuits. Indeed, previous reports have identified reduced functional connectivity in frontal regions of adult Shank3 mutant mice^52^, though it remains unclear whether this hypoconnectivity arises during early development. Future studies employing region-and cell-type-specific tools, coupled with *in vivo* electrophysiological techniques offering greater temporal resolution, will be critical for uncovering how Shank3 deficiency disrupts early activity patterns and different afferent input properties in L2/3 ACC circuits.

## Synaptic and Neuronal Adaptations in Adulthood

A striking aspect of our findings is the extensive neuronal adaptations observed in adulthood compared to early postnatal development. Most notably, PVIN excitability underwent a complete reversal, and in contrast to the hypoexcitability observed at P15, adult PVINs exhibited a significantly increased input-output I-F relationship in response to depolarizing current injections (Fig. 6I). Despite this enhanced excitability, glutamatergic input onto PVINs was markedly diminished, as reflected by a significant reduction in both mEPSC frequency and amplitude (Fig. 6C and 6D). Interestingly, optogenetically evoked IPSCs in PYRs following PVIN activation were not significantly different between genotypes (Fig. 6N), suggesting that total PVIN inhibitory output onto single PYR might be normalized over development despite extensive changes in PVIN synaptic and intrinsic properties. In parallel, PYR in layer 2/3 of the ACC also exhibited marked alterations in both intrinsic and synaptic properties by adulthood (Fig. 5). PYR excitability was significantly reduced, as indicated by a decreased I-F curve and prolonged AHP duration, phenotypes that may further reflect impairments in voltage-dependent potassium channels involved in AP repolarization and regulation of the slow components of AHP. Additionally, AMPAR-mediated mEPSCs exhibited reduced amplitude and slower kinetics, suggesting postsynaptic changes in PYR AMPAR properties. This depression of AMPAR synapses is consistent with a previous study that reported reduced mEPSC frequency and amplitude in L2/3 adult PYR^53^. However, we did not observe a significant difference in mEPSC frequency, which may be attributed to the higher mEPSC amplitude cutoff (7 pA) we used in our study. Together, these findings reveal an overall reduction in excitatory synaptic input onto both PVINs and PYRs in the ACC by adulthood, consistent with a broader weakening of glutamatergic connectivity in this region. The divergent changes in excitability, hyperexcitability in PVINs and hypoexcitability in PYRs, may reflect compensatory adaptations aimed at stabilizing circuit output in response to early inhibitory deficits. However, the extent to which these alterations restore normal network function remains unclear. It is also unknown whether these changes in PVIN and PYR phenotypes occur independently or through reciprocal interactions, highlighting the need for future studies dissecting the dynamic interplay between these neuronal populations throughout development. Importantly, our results demonstrate that the adult phenotypes in ACC of Shank3B^-/-^ mice do not simply represent persistent early deficits but instead reflect a complex trajectory of circuit reorganization. This underscores the importance of identifying and targeting pathophysiological processes during early postnatal development, before widespread compensatory changes obscure primary disease mechanisms. Such early-stage interventions may offer the greatest potential for restoring typical developmental trajectories in ASD and related neurodevelopmental disorders.

## Role of Shank3 in the postnatal maturation of cortical neurons

Shank3 is widely recognized as a postsynaptic scaffold protein at glutamatergic synapses^44^. However, our findings reveal that during early postnatal development, glutamatergic input onto both PYR and PVIN in layer 2/3 of the ACC remains largely intact in Shank3B^-/-^ mice (Fig. 1 and 2). This suggests that Shank3 is either dispensable or functionally compensated during the early stages of postnatal maturation of glutamatergic synapses *in vivo*. In contrast, PVIN excitability and HCN channel-mediated I_h_ currents are already disrupted at this stage (Fig. 2 and 4), pointing to impairments in non-synaptic mechanisms regulated by Shank3 that influence neuronal function. Interestingly, I_h_ currents in cortical PVIN are larger during early postnatal development, suggesting an important developmental role in PVIN maturation^67^. Previous studies have reported direct interaction between Shank3 and HCN channels^45,46^. However, the subcellular distribution of HCN channels differs markedly between PYR and PVIN, being primarily dendritic in PYR but enriched at the soma and axon initial segment in PVIN^78^. While some evidence from dissociated neuronal cultures suggests that Shank3 may be expressed in axons^85^, it remains unclear whether this is also observed *in vivo*, and whether axonal HCN expression can be directly regulated by Shank3. As an alternative to direct protein-protein interaction, Shank3 may also influence mechanisms controlling neuronal excitability through transcriptional regulation. Shank3 has been reported to translocate to the nucleus^86^ and indirectly modulate nuclear signaling via calcium-dependent pathways mediated by L-type voltage-gated Ca²⁺ channels^86–88^. Given the crucial role of calcium dynamics and buffering in PVIN physiology, including regulation of downstream transcriptional regulation, PVINs may be especially vulnerable to disrupted calcium homeostasis following Shank3 loss. Although the precise mechanisms by which Shank3 influences PVIN excitability and axonal function remain to be determined, our data support a model in which non-synaptic roles of Shank3, particularly in regulating HCN channel function and AP firing, play a critical role in postnatal cortical development and should be considered when interpreting neuronal and circuit level phenotypes associated with SHANK3 dysfunction.

## Implications for ASD research

Individuals with SHANK3-related neurodevelopmental disorders frequently present with seizures and exhibit epileptiform activity in EEG recordings^9,42,89^. Our findings reveal that PVIN dysfunction emerges early in development, preceding more pronounced synaptic deficits observed in adulthood. This temporal sequence suggests that early interventions aimed at restoring PVIN excitability may represent a promising strategy for preventing cortical activity imbalances in SHANK3-associated conditions. In addition, PVINs in the human mPFC exhibit significantly larger I_h_ currents compared to those in rodents^90,91^, implying that HCN channel dysfunction may have a more profound impact in the human brain. Importantly, enhancing GABAergic function has shown efficacy in rescuing abnormal behaviors in several mouse models carrying ASD-linked mutations^91–94^. Our study provides new insights into the developmental trajectory of PVIN and PYR neuron dysfunction in the ACC of Shank3B^-/-^ mice and reinforces the notion that early PVIN dysfunction is a key driver of cortical imbalances in ASD.

## Acknowledgements

We thank Tasha Merchant, Lan Chen and Sidney Dawkins for assistance with mouse husbandry and genotyping. We thank Susana da Silva, David Lewis, Guillermo Gonzalez-Burgos, Gary Thomas and Stephanie Rudolph for helpful discussions and comments on the manuscript. R.T.P was supported by R01MH124695, R21MH132015 and a Bridge to Independent Award from the Simons Foundation Autism Research Initiative. L.N. was supported by T32MH16804.

## Author Contributions

R.T.P. conceived and designed the study. Y-C.S., L.N., M.J., M.M. and A.D’A. performed electrophysiological experiments, and Y-C.S. analyzed the electrophysiological data. Y-C.S. collected and analyzed the two-photon microscopy data. Y-C.S. and M.M. performed immunohistochemistry. L.N. developed scripts for electrophysiological data analysis, and Y-C.S. and R.T.P. developed scripts for two-photon data analysis. Y-C.S. and R.T.P. wrote the manuscript with input from all the other authors.

## Conflict of Interest Statement

The authors declare that they have no conflicts of interest related to the content of this manuscript. No author has any financial or personal relationships with individuals or organizations that could inappropriately influence, or be perceived to influence, the research and findings reported in this paper. Additionally, the authors have no competing financial interests related to the content of this manuscript.

## Methods Animals

All experimental manipulations on mice were performed in accordance with protocols approved by the Institutional Animal Use and Care Committee at the University of Pittsburgh in compliance with the guidelines described in the US National Institutes of Health *Guide for the Care and Use of Laboratory Animals*. Mice were housed on a 12/12hr light/dark cycle with chow and water provided ad libitum. Mice were weaned at P21-23 and separated by sex in cages of 2-5 animals of mixed genotypes. Shank3B^-/-^ mutant mice were described previously^61^ and obtained from The Jackson Laboratory (#017688). Characterization of neural properties by electrophysiology was performed in Shank3B^+/+^ and Shank3B^-/-^ age-matched mice from breeding pairs between Shank3B^+/-^ heterozygous animals. Male mice were used in all experiments. Animals were excluded from experiments if their bodyweight was smaller than 4.5g at P15, or substantially smaller than their littermates. Due to the low yield of our genetic crosses and the requirement for genotyping prior to experimentation, randomization and blinding of sample analysis were not feasible.

## Viruses and stereotaxic intracranial injections

Intracranial viral injections were performed at P7-P8 (for experiments performed at P15) or 2 weeks before experiments (for experiments performed in adult mice). Mice were anesthetized with isoflurane (4% for induction and 1-2% thereafter for maintenance) and placed into a stereotaxic apparatus (Kopf Instruments model 1900). The hair around the site of incision was removed using a depilatory cream (Nair). The surgical area was cleaned by 2% iodine and 70% alcohol solution. To perform intracranial injections, injection sites were poked by sterile 30G needle in P7-P8 pups and drilled by Micromotor drill (Stoelting) in adults, respectively. All skull measurements were made relative to Bregma, and AAV solutions were delivered by injecting 350 nl at a maximum rate of 75 nL/min using a Harvard Apparatus PHD ULTRA CP injector. For labeling of ACC PVIN, an adenoassociated virus (AAV) expressing tdTomato under the PVIN specific S5E2 enhancer (AAV9-S5E2-tdTomato, Addgene 135630-AAV9) was injected in mice using coordinates: AP +0.5 mm; ML 0.3 mm; Depth -1.0 mm for P7 pups and AP +1.1 mm; ML 0.35 mm; Depth -1.1 mm for adults. AAV9-S5E2-C1V1-YFP (Addgene 135633-AAV9) was used for expressing the red-shifted Channelrhodopsin C1V1 in PVIN. Co-expression of GCaMP8s and S5E2-tdTom in ACC for two-photon imaging experiments was achieved by injecting AAV-syn-jGCaMP8s-WPRE (Addgene 162374-AAV9) and AAV9-S5E2-tdTomato at a 2:1 ratio. AAV-hChR2(H134R)-mCherry (Addgene 26976-AAV8) was used for expressing ChR2 in MD with coordinates: AP -0.9 mm; ML 0.4 mm; Depth -3.2 mm for P7 pups. Mice received carprofen (5 mg/kg) for analgesia before surgery. Following injections and wound suture, mice were placed on an heating pad until being fully awake and returned to their home cages for at least 7 days.

## Brain slice preparation and whole-cell electrophysiology

Acute brain slices were prepared following anesthesia by isoflurane inhalation and transcardiac perfusion with ice-cold artificial cerebrospinal fluid (ACSF) containing (in mM): 125 NaCl, 2.5 KCl, 25 NaHCO_3_, 2 CaCl_2_, 1 MgCl_2_, 1.25 NaH_2_PO_4_ and 25 glucose (310 mOsm per kg). Cerebral hemispheres were removed and transferred into a slicing chamber containing ice-cold ACSF. Coronal slices including ACC (275 µm thick) were cut with a Leica VT1200s vibratome and transferred for 10 min to a holding chamber containing choline-based solution consisting of (in mM): 110 choline chloride, 25 NaHCO_3_, 2.5 KCl, 7 MgCl_2_, 0.5 CaCl_2_, 1.25 NaH_2_PO_4_, 25 glucose, 11.6 ascorbic acid, and 3.1 pyruvic acid at 33°C. Slices were subsequently transferred to a chamber with pre-warmed ACSF (33°C) and gradually cooled down to room temperature (20–22°C). All recordings were obtained within 4 hours of slicing. Both ACSF and choline solution were constantly bubbled with 95% O_2_ and 5% CO_2_. Individual slices were transferred to a recording chamber mounted on an upright microscope (Olympus BX51WI) and continuously perfused (1–2 ml per minute) with ACSF at room temperature. Cells were visualized using a 40× water-immersion objective with infrared illumination. Whole-cell voltage clamp recordings were made from PYR or PVIN of ACC. Patch pipettes (2.5–4 MΩ) pulled from borosilicate glass (BF150-86-7.5, Sutter Instruments) were filled with a Cs^+^-based internal solution containing (in mM): 130 CsMeSO_4_, 10 HEPES, 1.8 MgCl_2_, 4 Na_2_ATP, 0.3 NaGTP, and 8 Na_2_-phosphocreatine,10 CsCl_2,_ 3.3 QX-314 (Cl^−^ salt), (pH 7.3 adjusted with CsOH; 300 mOsm per kg) for voltage-clamp recordings. Alexa Fluor 488 dye (0.07%) was added to the internal solution to fill the patched cell and examine dendritic morphology. PYR identity was validated by the presence of an apical dendrite and spiny dendrites, whereas PVIN were identified by S5E2-Tom expression and validated by aspiny dendritic morphology. For all voltage-clamp experiments, errors due to voltage drop across the series resistance (<20 MΩ) were left uncompensated. For mEPSC recordings, ACSF contained 1 μM TTX, 1 μM (RS)-CPP, and 1 μM Gabazine, and recordings were performed at room temperature (20-22°C) with V_m_ = -70 mV. After breaking in, cells were left to stabilize for 4 min and currents were then acquired continuously for 5 min. Membrane currents and potentials were amplified and low-pass filtered at 3 kHz using Multiclamp 700B amplifier (Molecular Devices), digitized at 10 kHz, and acquired using National Instruments acquisition boards and a custom version of ScanImage written in MATLAB (Mathworks). Calculation of input resistance and membrane capacitance in voltage clamp recordings was performed by fitting evoked currents in response to -5 mV voltage steps in the first seconds after cell break-in. For HCN I_h_ current isolation, ACSF contained 1 μM TTX, 100 μM 4-AP, and 1 mM BaCl_2_, and recordings were performed at room temperature with Vm = -60 mV. Potassium-based internal solution containing (in mM): 120 KMeSO_3_, 10 HEPES, 20 KCl, 0.1 EGTA, 4 Na2ATP, 0.3 NaGTP, 5 Na_2_-phosphocreatine, and 30 tetraethylammonium (pH 7.3 adjusted with KOH; 310 mOsm per kg) was used. Cell membrane potential was changed with a series of hyperpolarized voltage steps (-10 to -100 mV in -10 mV increments) for 2s. HCN channel blocker, ZD7288 (Tocris, 50 μM), was added into both ACSF and internal solution for confirming HCN current. For current-clamp recordings, potassium-based internal solution containing (in mM): 130 KMeSO_3_, 10 HEPES, 3 KCl, 1 EGTA, 4 Na_2_ATP, 0.3 NaGTP, and 8 Na_2_-phosphocreatine, (pH 7.3 adjusted with KOH; 300 mOsm per kg) was used. No drug was added to the bath, and recordings were performed at 31-33°C. Cell was broken in with V_m_ = -70 mV and allowed to stabilize for 4 min in voltage-clamp. After stabilization, resting membrane potential was measured at I = 0. Current-clamp recording was performed by adjusting holding current to maintain the V_m_ at -70mV. Four cycles of 700ms current steps -100 pA to 350 pA in 25 pA increments were injected and V_m_ changes recorded and averaged. For oIPSC recordings, voltage-clamp was performed in Cs^+^-based internal solution for measuring optogenetically evoked synaptic currents. Cells were held at V_m_ = 0 mV in ACSF with 1 μM (RS)-CPP and 1 μM NBQX. C1V1-dependent oIPSC was induced by a 1ms 5.5mW 563nm light pulse (CoolLED pE-300) delivered through the objective for 4-5 cycles (45s interval). For controlling fiber density, YFP fluorescence in the field of view was imaged with 10.2 mW 473nm light for 100ms, centering on patched cell. For controlling the efficacy of opsin activation between genotypes, success rate of AP and opsin-dependent current in C1V1-expressing cells were characterized by current clamp and voltage clamp, respectively. Standard potassium-based and Cs-based internal solutions were used as described above. ACSF contained 1 μM (RS)-CPP, 1 μM NBQX, and 1 μM Gabazine, and recordings were performed at room temperature with Vm = -70 mV. C1V1 was activated with the same light stimulation condition used for oIPSC PYR recordings, and YFP fluorescence of cell body was quantified for oEPSC amplitude normalization. For recordings of MD-driven feedforward inhibition, 1 μM (RS)-CPP was added to the ACSF, and a modified low-chloride Cs^+^-based internal solution containing (in mM): 130 CsMeSO_4_, 10 HEPES, 1 EGTA, 4 Na_2_ATP, 0.3 NaGTP, and 8 Na_2_-phosphocreatine, 3.3 QX-314 (Cl^−^ salt) was used. Cells were held at -70 mV and 0 mV for measuring AMPA and GABA currents, respectively. Thalamocortical ChR2-dependent oEPSC was induced by 1ms 7mW 473nm light pulse. mCherry intensity in the field of view was captured with 5.5mW 563nm LED for 100ms. For comparing NMDAR / AMPAR (N/A) cureent ratio, MD-driven AMPAR-dependent and NMDAR-dependent oEPSC was measured by holding cells at -70 mV and +40 mV, respectively. ACSF contained 1 μM Gabazine, and voltage clamp recordings were performed at room temperature. ChR2-dependent oEPSC was induced by 1ms 473nm light pulse and the light power was tuned for each cell to induce an AMPAR-dependent current in the range between 100 to 700 pA for prevent unclamped action potentials and minimize nonlinearities caused by high currents. Electrophysiology recordings were performed in a custom version of ScanImage and saved as Matlab files for subsequent off-line analysis using the custom Python program ClampSuite.

## Brain tissue imaging and immunohistochemistry

Mice were deeply anesthetized with isofluorane and perfused transcardially with 1X phosphate-buffered saline (PBS) (pH 7.4) followed by 4% paraformaldehyde in PBS. Brains were fixed overnight at 4°C, washed in phosphate buffer saline (PBS) and sectioned (50 µm) coronally using a vibratome (Leica VT1000s). For immunohistochemistry, brain sections were incubated in blocking buffer containing: 10% normal goat serum, mouse IgG (1 μg/mL) and 0.3% triton X-100 in 1X PBS for one hour at room temperature. After blocking, sections were incubated overnight with primary antibodies: anti-Parvalbumin (1:10000, SWant PV235), anti-VVA biotinylated (1:400, Vector Laboratories), and anti-Somatostatin (1:500, Millipore Sigma MAB354) at 4°C. After the sections were washed, they were incubated with anti-rat Alexa 488 (1:2000, Jackson ImmunoResearch), anti-mouse Alexa 647 (1:2000, Abcam), Streptavidin-Alexa 594 (1:2000, Jackson ImmunoResearch), and DAPI (1 μg/mL, Thermo Fisher) for 1-2 hours at room temperature. The sections were mounted on glass slides, dried and mounted with ProLong antifade reagent containing DAPI (Molecular Probes). Whole brain sections were imaged using and Olympus VS200 slide scanner with 10X and 40X objective lens. For quantification of PV-tdTom co-localization slices were imaged in an Olympus FV1200 confocal microscope with 60X objective lens. Three slices containing ACC were analyzed per animal to quantify percentage of PV-positive cells among S5E2-tdTom labeled population. For each slice three regions of L2/3 were randomly selected and imaged, and cell number was manually quantified. PV signal was analyzed in ImageJ and considered positive if fluorescence intensity was 2.5x above the background. A population of S5E2-tdTom-labeled cells with very small cell size (<100 μm^2^) in adult ACC was excluded from quantification and whole-cell patch clamp recordings. For quantification of PV immunoreactivity across layer 2/3 and layer 5 of ACC and S1 we imaged four coronal brain sections per animal. Sections were selected from alternating hemispheres and PV signal was analyzed separately for each cortical region and layer. Separate masks were defined for each layer and cortical region and further analyzed for cell segmentation. For each layer mask we quantified three user-defined background ROIs for signal subtraction and calculated the total area for cell density normalization. PV+ cell detection was performed using CellPose with a fixed segmentation diameter of 33 pixels on the fluorescence channel corresponding to PV staining. Segmentation results were manually curated to remove misidentified ROIs, and the final outlines were exported to ImageJ for analyzed of fluorescence intensity of cell segmentation ROIs. Cell density was calculated by dividing the total PV+ cell count by the measured area of the region.

## Cranial window implants

P15-P17 mice previously co-injected with AAV-GCaMP8s and AAV-S5E2-tdTom were anesthetized with isoflurane as previous described. Mice received carprofen for analgesia before surgery. Eye ointment (Systane) was applied to prevent eyes dry during the surgery. The hair on scalp was removed using the hair removal cream (Nair) and the surgical area further cleaned by 2% iodine and 70% ethanol solution. The scalp over the surgical area was removed and the skull was cleaned again with 2% iodine and 70% ethanol solution. The skull above the injection site was carefully drilled using a micromotor drill to open a craniotomy of approximately 3.5 mm diameter. The drilling site was rinsed with ice-cold sterile saline after every round of drilling. Ice-cold 0.01% epinephrine solution (Fisher scientific) was injected under the skull, and bone fragments were carefully removed with sharp forceps. The dura was kept intact and moist with sterile saline. The exposed area of the brain was routinely rinsed with ice-cold epinephrine solution until no bleeding was observed. Pre-sterilized 5 mm diameter coverslips (Thermofisher) were then gently placed on the brain surface with sterile saline added between the coverslip and the brain. The edges of the coverslip were glued to the skull using LOCTITE 454 superglue, and a custom titanium headpost was subsequently secured using Metabond cement (C&B Metabond). Following surgery, mice were placed on a heating pad until awakening. Grated rimadyl (Bio-Serv, 2 mg/tb) mixed with RO water was provided during recovery time before recordings.

## Two-photon calcium imaging

Two hours following recovery from window implantation, mice were head-fixed on top of a heating pad (37°C) in a custom stage with goniometers for fine tilt adjustments. Images were acquired using a resonant scanning two-photon microscope (Ultima 2P plus, Bruker, WI) at ∼15 Hz frame rate and 512x512 pixel resolution through a 16x water immersion lens (Nikon 16x 0.8NA). Dorsal ACC was imaged at a depth between 150 and 350 μm with GCaMP8s and S5E2-tdTom expression.

Excitation light (920 nm/1100nm wavelength) was provided by a tunable femtosecond infrared laser (Insight X3, Spectra-Physics, CA). Tunable wavelength beams were combined with a dichroic mirror (ZT1040dcrb-UF3, Chroma, VT) before being routed to the microscope’s galvanometers. PrairieView software (vX5.5 Bruker, WI) was used to control the microscope. Multiple 6 min imaging (5380 images at ∼15 Hz frame rate). were acquired from each mouse at varying depths. After each imaging session, 20 μm z-stacks with 5 μm spacing were imaged with 1100 nm excitation for S5E2-tdTom visualization. Mediolateral distance of the field of view was measured in relation to the central sinus to estimate cortical layer.

## Two-photon calcium imaging data analysis

Sequential images of GCaMP8s fluorescence were registered and motion corrected in Suite2p (v0.10.3, Cellpose v0.7.3). Regions of interest (ROIs) were identified based on functional segmentation by correlating pixel intensity across time. Z-stack images of S5E2-tdTom fluorescence collected with 1100 nm laser excitation were max projected into a single plane in ImageJ, and S5E2-tdTom positive cells were identified using CellPose (Strubger et al. 2021). Data was further processed using custom Python scripts for movement artifact removal, signal deconvolution, and Ca^2+^ transient quantification. No randomization or blinding of analysis were performed. Movement artifacts were identified by calculating the spatial displacement of the imaging plane across frames using the vectorized xoff/yoff shift from the Suite2p ops.npy file. Image segments with frame-to-frame shifts exceeding 1.5 pixels were removed and surrounding frames were concatenated and smoothed using a Savitzky-Golay filter (window size of 15 frames, polynomial order of 3). Fluorescence traces were corrected for neuropil contamination by subtracting a scaled version of the neuropil signal (F_subt = F -0.7*Fneu). Baseline fluorescence (F0) was estimated using a rolling 8th percentile function over a 120-second window^107^. The normalized change in fluorescence (ΔF/F) was calculated as ΔF/F = (F − F0) / F0. For Ca^2+^ event quantification, we detected peaks in the filtered ΔF/F trace using the SciPy findpeaks function with the following parameters: [PYR: distance = 3, prominence =0.5, width = 3; PVIN: distance = 2, prominence =0.4, width = 1]. Event amplitude, frequency, area under the curve (AUC), rise time, tau, and duration were calculated to quantify calcium event properties. To account for phase shifts introduced by wavelet filtering, each detected peak position was redefined by searching for the local maximum in the unfiltered ΔF/F trace within 3 frames around the identified peak. To determine the onset (and thereby the baseline) of an event, we first identified the minimum ΔF/F value in the 20 frames preceding the event peak (equivalent to 1.33 seconds at 15 Hz). For events with ΔF/F values below 1.5, we reduced the search window to 15 frames. From this minimum, we considered all indices where ΔF/F values were less than the minimum + 0.15. We then calculated the slopes from these points to the event peak, identifying potential onset indices as those with a slope greater than max slope -0.03. Among these candidates, we selected the index closest to the peak as the event onset. Event amplitude was calculated as the ΔF/F value at peak minus the value at peak onset. The end of the event was defined as the point at which ΔF/F reached 37% of peak amplitude, or the onset of the next peak, whichever happened first. Peak area under the curve (AUC) was measured by integrating the values from the event onset to event end. ROIs were excluded based on the following criteria: [all event parameters (frequency, amplitude, etc) = 0, noise size > 1, F < 80, and F_subt < 40]. For activity correlation analysis, we used deconvolved data to avoid overestimation of correlation between ROIs. Fluorescence traces were deconvolved using the OASIS AR(1) model with FOOPSI and only deconvolved events with amplitudes exceeding 0.05 were retained for further analysis. Deconvolved traces were then smoothed using a 300 ms moving average and further used to compute the correlation between neural activity. Euclidian distances between cells were calculated as xy vector length between each ROI centroid. We calculated pairwise Pearson correlations between all ROI pairs from each recording and generated correlation and distance matrices. The averaged cumulative distribution of Pearson correlations was measured by averaging the cumulative distribution of all Pearson coefficients of each individual animal. These analyses were performed separately for both PYR and PVIN cell populations.

## Electrophysiology data analysis

mEPSCs, current-clamp, and oEPSC/oIPSC were analyzed using a custom python-based program Clampsuite. Acquisitions had the mean subtracted and were filtered using a zero-phase Remez filter with a low pass filter at 600 Hz and 600 Hz wide. Events were identified using a modified FFT deconvolution method. Events were discarded based on the following criteria: Events smaller than 7 pA (measured from the baseline to the peak), a rise time faster than 0.5 ms, a rise time (time from baseline to event peak) slower than the decay, a rise time slower than 4 ms, a minimum decay time of 0.5 ms and a minimum interevent distance of 1 ms. Tau was estimated as the time when the event reached 37% of peak amplitude. Rise time was calculated as the time from the start of the mEPSC to the peak. Rise rate is the amplitude of the divided by the rise time. Occasionally an event was manually removed or labeled. For analysis of optogenetically evoked currents, oEPSC peak was identified in the 5-20 ms poststimulation window, and oIPSC peak was identified 5-50 ms after stimulation. For N/A ratio measurement, AMPAR current amplitude at -70 mV was identified in the 5-20 ms poststimulation window; NMDAR current amplitude at +40 mV was identified at 50 ms poststimulation. For analysis of current clamp recordings, evoked potentials were analyzed in ClampSuite. All AP statistics, except AP firing rate, were calculated from the first acquisition of each cycle (ranging from –100 to 300 pA in order from most negative to most positive) that contained at least one AP and taking the mean of the values of all APs in that cycle. The following were calculated from the first AP of each acquisition containing a spike: AP threshold, peak AP velocity, start of the AHP, AP width, AP peak voltage, and AP duration. The AP threshold was identified by analyzing the third derivative. The peak AP velocity was the peak of the first derivative of the AP. The start of the AHP was calculated from the third derivative of the AP. The AP peak voltage was calculated as the maximum voltage of the AP. The membrane resistance was calculated from the first 7 pulses (-100 to 50 pA) by regressing the ΔV (y) against the current pulse amplitude (x) and taking the slope as the membrane resistance. Vm was calculated as the mode of the moving average during the step duration. The AP frequency (Hz) of each pulse step was the average number of spikes divided by the time of each pulse step. Acquisitions without spikes were assigned a null value and did not count towards the mean frequency for each pulse amplitude. For I_h_ current isolation, the trace was denoised to remove mEPSCs. The peak current was identified in 100-300 ms post-hyperpolarized step window, and the baseline was identified in 1-2 s and 0.35-1 s post-hyperpolarized step window for PYR and PVIN, respectively. For decay tau measurements, I_h_ current traces were fit with a single-order exponential. For electrophysiology data analysis, no sample randomization or blinding were used.

## Statistical analyses

Sample sizes were selected based on prior similar studies and expected effect sizes. We tested the normality of all collected datasets using QQ plots. Since not all the datasets were normally distributed, hypothesis testing between two groups was performed using the Mann–Whitney U (MW) test. Outlier mice were excluded if the primary measurement fell outside the range of mean ± 2 standard deviations. If the average value for an animal met this criterion, all samples from that animal were excluded from analysis. Cumulative distribution functions (CDFs) were measured for individual samples and reshaped to equal sample size (n = 500), and the empirical CDF (eCDF) was calculated by averaging those CDFs. The Kolmogorov-Smirnov (KS) test was employed to compare eCDFs between groups. Two-way ANOVA was performed for comparing parameters affected by two factors. Sigmoid curve fit was performed for estimating the max current density (A) and slope (k) of V_m_-I_h_ density curves. Values are presented as median (center line), with the 25th (lower bound) and 75th percentiles (upper bound) represented by the box, and whiskers indicating the range in box plots. Statistical significance was considered at P < 0.05. Statistical calculations were performed using SciPy (Python). Specific statistical analyses are detailed in the respective figure legends or results section for each dataset.

## Code availability

Python scripts used for the analysis of two-photon calcium imaging and whole-cell electrophysiology data are available at https://github.com/Peixoto-Lab/Shih_2025. ClampSuite used for processing raw electrophysiological recordings is available at https://github.com/LarsHenrikNelson/ClampSuite.

---

**Supplementary figure 1.**
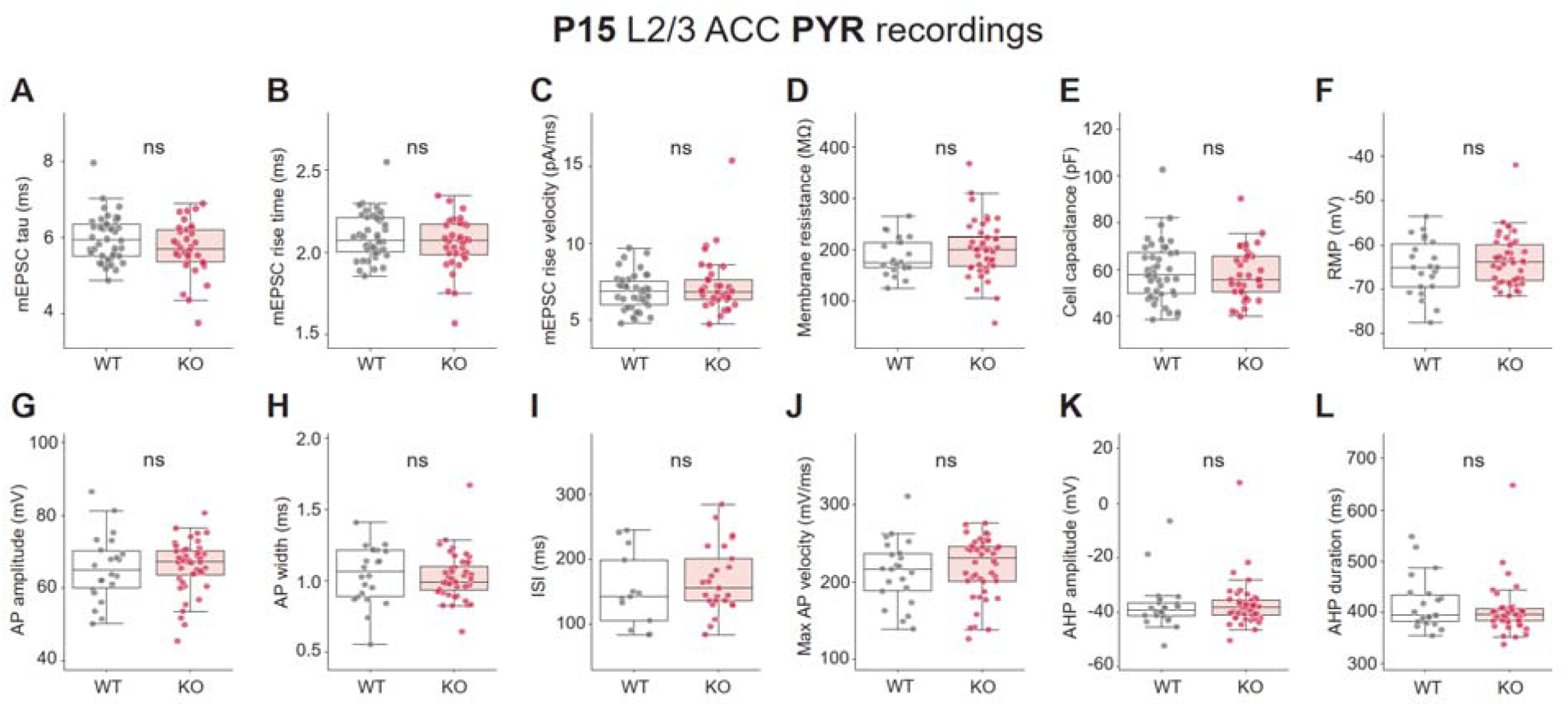
mEPSC and action potential properties of L2/3 PYR in ACC of P15 Shank3B^-/-^ mice. **(A)** mEPSC tau of PYR (Average±SD, WT 6.0±0.6ms; KO 5.7±0.7ms, MW test p=0.17). **(B)** mEPSC rise time of PYR (Average±SD, WT 2.1±0.1ms; KO 2.1±0.2ms, MW test p=0.37). **(C)** mEPSC rise velocity of PYR (Average±SD, WT 6.8±1.2pA/ms; KO 7.3±1.9pA/ms, MW test p=0.6). **(D)** Membrane resistance of PYR (Average±SD, WT 186.9±36.5MΩ; KO 200.7±54.8MΩ, MW test p=0.28). **(E)** Cell capacitance of PYR (Average±SD, WT 59.1±12.9pF; KO 57.9±11.4pF, MW test p=0.65). **(F)** Resting membrane potential of PYR (Average±SD, WT - 65.0±6.3mV; KO -63.5±5.7mV, MW test p=0.47. **(G)** AP amplitude of PYR (Average±SD, WT 65.5±9.2mV; KO 66.0±7.2mV, MW test p=0.58). **(H)** AP width of PYR (Average±SD, WT 1.0±0.2ms; KO 1.0±0.2ms, MW test p=0.48). **(I)** Inter-spike interval of PYR (Average±SD, WT 153.1±57.7ms; KO 168.6±51.8ms, MW test p=0.44). **(J)** Maximum AP velocity of PYR (Average±SD, WT 213.0±41.7mV/ms; KO 219.3±38.9mV/ms, MW test p=0.38). **(K)** Afterhyperpolarization amplitude of PYR (Average±SD, WT - 37.5±9.2mV; KO -37.1±8.7mV, MW test p=0.57). **(L)** Afterhyperpolarization duration of PYR (Average±SD, WT 416.3±51.4ms; KO 403.0±48.0ms, MW test p=0.51). For mEPSC recordings, n=42,35 neurons from n=4,3 WT and Shank3B^-/-^ mice. For current-clamp recordings, n=25,44 neurons from n=4,5 WT and Shank3B^-/-^ mice. Box plots show median and quartile values. Box plots *P<0.05 with MW test.

**Supplementary figure 2.**
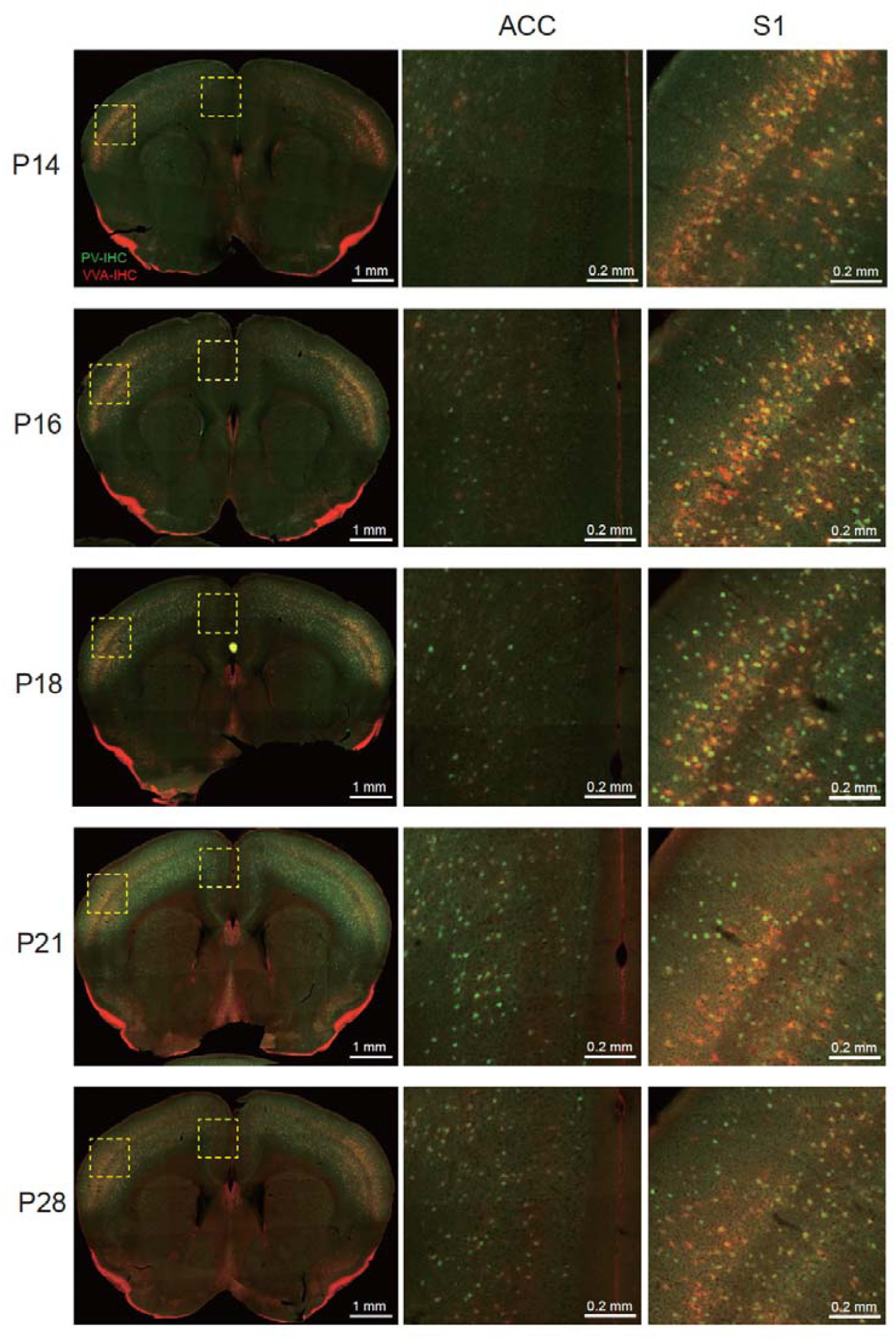
Developmental expression of Parvalbumin and VVA in the anterior cingulate cortex (ACC) and somatosensory cortex (S1). Representative coronal brain sections of WT mice at postnatal days P14, P16, P18, P21, and P28. Immunohistochemistry for Parvalbumin (PV, green) and the perineuronal net marker Vicia Villosa Agglutinin (VVA, red) shows the distribution and density of PV-INs and VVA nets in the ACC (middle panels) and S1 (right panels). The yellow dashed boxes in the whole brain images (left panels) indicate the regions of the ACC and S1 that are magnified in the middle and right columns, respectively. Note that expression of PV in ACC is delayed in comparison with S1, only reaching full uniform expression by P21. Scale bars: 1 mm for whole brain sections, 0.2 mm for magnified regions.

**Supplementary figure 3.**
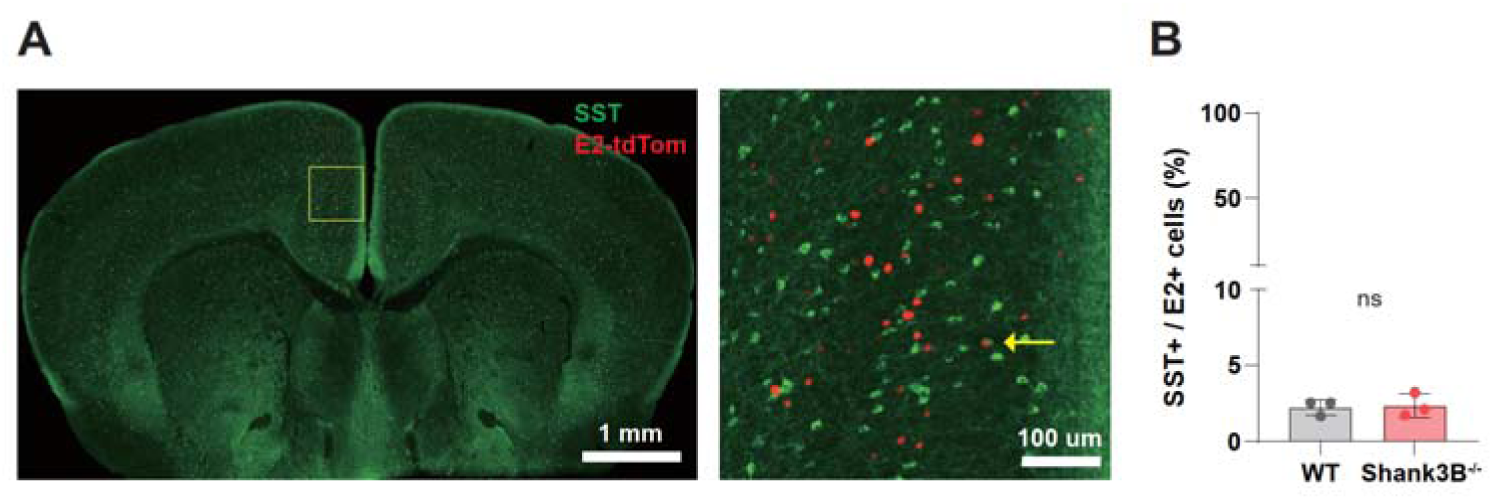
Percentage of S5E2-Tom positive interneurons overlapping with Somatostatin IHC in L2/3 ACC of P15 WT and Shank3B^-/-^ mice. (A) Representative coronal brain section of WT mice at postnatal days P15. Immunohistochemistry for somatostatin (SST, green) and S5E2-tdTom positive neurons in ACC. The yellow arrow in magnified image (right panel) indicates the S5E2-tdTom^+^ cell overlapping with SST. (B) Average percentage of S5E2-tdTom^+^ cells overlapping with somatostatin signals (Average±SD: WT 2.27±0.50%; KO 2.37±076%). Scale bars: 1 mm for whole brain sections, 0.1 mm for magnified region. n=3,3 WT and Shank3B^-/-^ mice.

**Supplementary figure 4.**
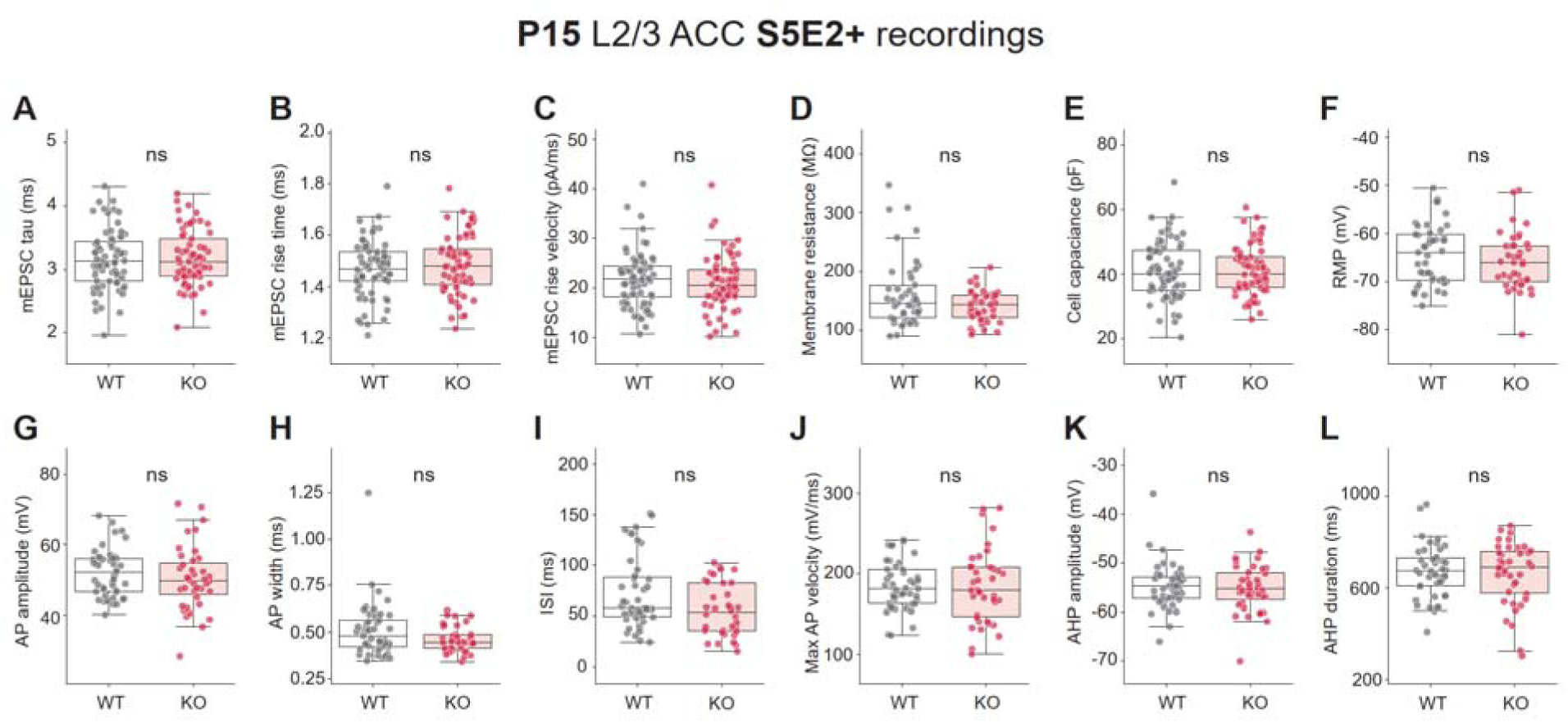
mEPSC and action potential properties of L2/3 PVIN in ACC of P15 Shank3B^-/-^ mice. **(A)** mEPSC tau of S5E2-Tom+ interneurons (Average±SD, WT 3.2±0.5ms; KO 3.2±0.4ms, MW test p=0.92). **(B)** mEPSC rise time of S5E2-Tom+ interneurons (Average±SD, WT 1.5±0.1ms; KO 1.5±0.1ms, MW test p=0.47). **(C)** mEPSC rise velocity of S5E2-Tom+ interneurons (Average±SD, WT 21.7±5.5pA/ms; KO 21.1±5.5pA/ms, MW test p=0.52). **(D)** Membrane resistance of S5E2-Tom+ interneurons (Average±SD, WT 162.1±58.3 MΩ; KO 140.7±27.5 MΩ, MW test p=0.25). **(E)** Cell capacitance of E2-Tom+ interneurons (Average±SD, WT 41.1±8.8pF; KO 41.2±7.6pF, MW test p=0.94). **(F)** Resting membrane potential of S5E2-Tom+ interneurons (Average±SD, WT - 64.3±6.2 mV; KO -65.9±5.9mV, MW test p=0.28). **(G)** AP amplitude of S5E2-Tom+ interneurons (Average±SD, WT 52.0±6.8 mV; KO 50.4±9.1 mV, MW test p=0.33). **(H)** AP width of S5E2-Tom+ interneurons (Average±SD, WT 0.5±0.2ms; KO 0.5±0.1ms, MW test p=0.21). **(I)** Inter-spike interval of S5E2-Tom+ interneurons (Average±SD, WT 72.7±35.4ms; KO 55.7±25.8ms, MW test p=0.21). **(J)** Maximum AP velocity of S5E2-Tom+ interneurons (Average±SD, WT 184.7±29.7mV/ms; KO 184.2±25.8mV/ms, MW test p=0.86). **(K)** Afterhyperpolarization amplitude of S5E2-Tom+ interneurons (Average±SD, WT -54.7±4.7mV; KO -55.2±4.7mV, MW test p=0.72). **(L)** Afterhyperpolarization duration of E2-Tom+ interneurons (Average±SD, WT 672.2±112.7ms; KO 662.2±136.1ms, MW test p=0.8. For mEPSC recordings, n=69,65 neurons from n=6,6 WT and Shank3B^-/-^ mice. For current-clamp recordings, n=46,39 neurons from n=4,4 WT and Shank3B^-/-^ mice. Box plots show median and quartile plots. Box plots *P<0.05 with MW test.

**Supplementary figure 5.**
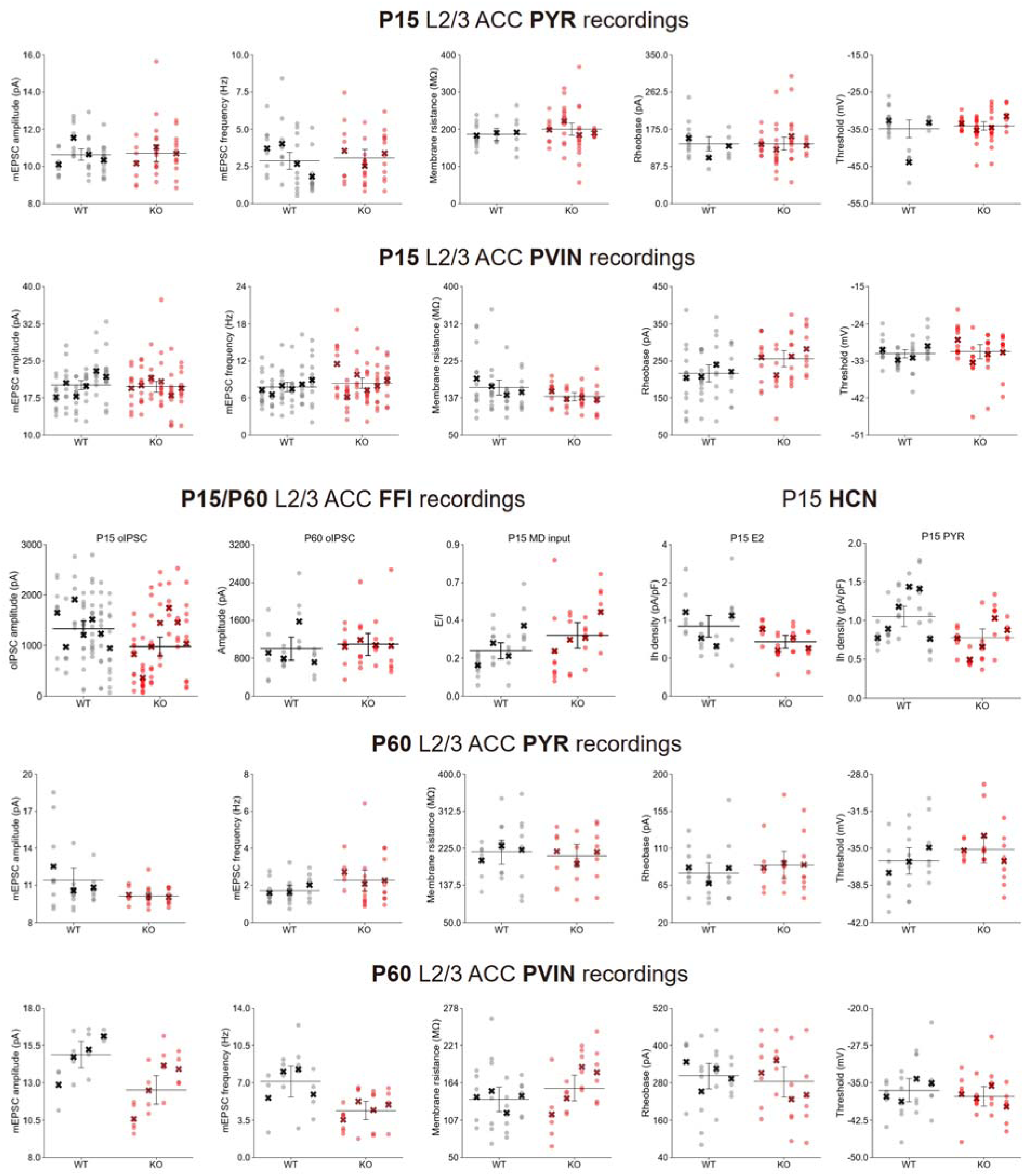
Mouse average and sample distribution of individual animals. Each plot shows average and SEM of mouse averages from the same genotype. Samples from the same animal are arranged in the same column. X marks the average of each neuron parameter for each mouse.

**Supplementary figure 6.**
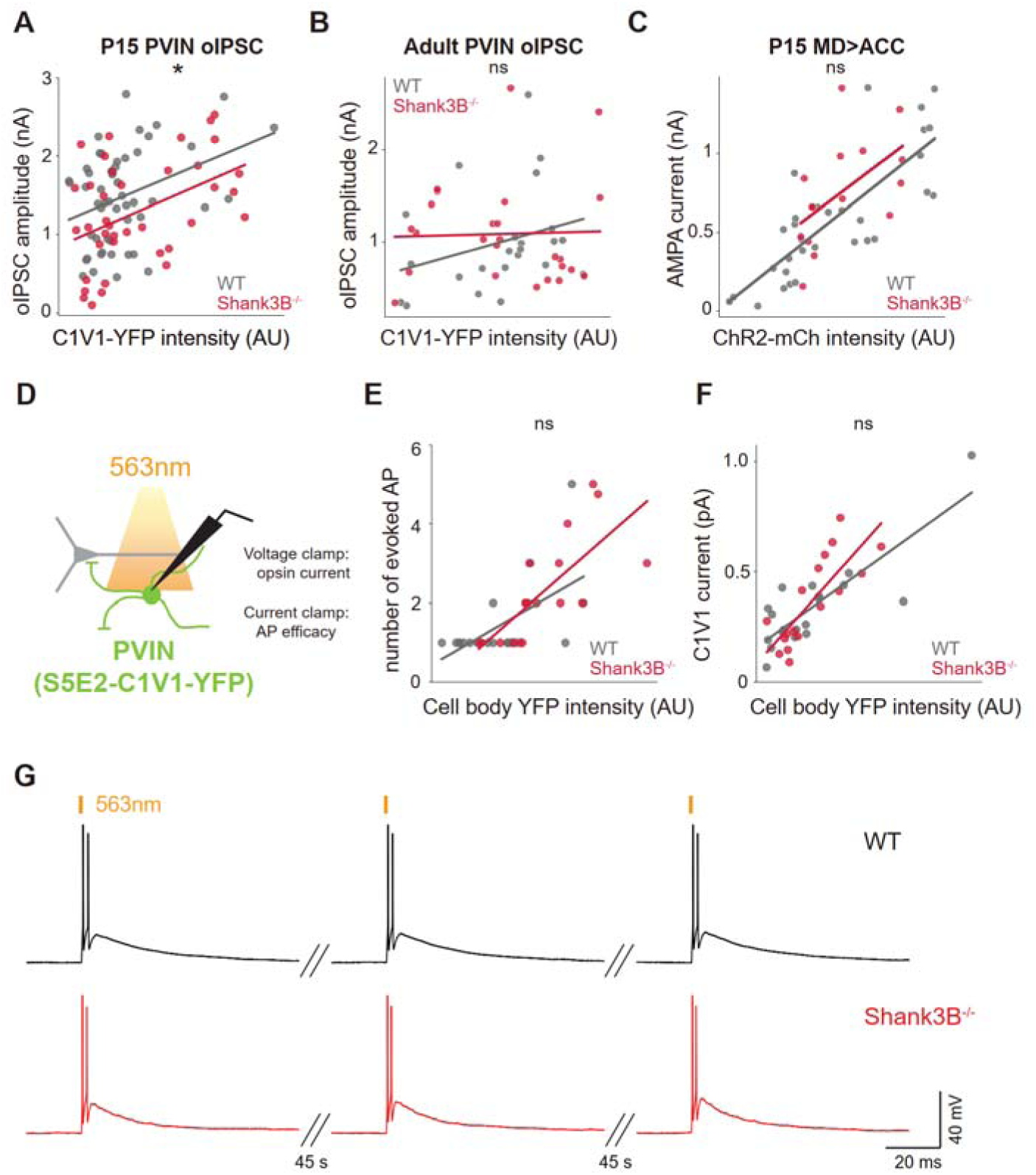
Relationship between opsin expressing fiber fluorescence intensity and oIPSCs and PVIN C1V1 efficacy control experiments. (A) Relationship between YFP intensity (C1V1-YFP expressing S5E2^+^ interneurons) and oIPSC amplitude evoked on L2/3 ACC PYR in P15 (Two-way ANOVA p=0.036 in genotype) and (B) adult WT (black) and Shank3B^-/-^ (red) mice (two-way ANOVA p = 0.33 in genotype). (C) Relationship between mCherry intensity (ChR2-mCh expressing fibers from MD) and AMPA currents evoked on L2/3 ACC PYR in P15, two-way ANOVA p=0.12 in genotype. (D) Schematic representing whole-cell recordings of S5E2-C1V1-YFP^+^ interneurons in L2/3 ACC of P15 mice. (E) Relationship between YFP intensity (cell body of C1V1-YFP expressing S5E2^+^ interneurons) and number of APs evoked in L2/3 ACC YFP^+^ interneurons in P15 (Two-way ANOVA p=0.93 in genotype) and (F) relationship between YFP intensity (cell body of C1V1-YFP expressing S5E2^+^ interneurons) and amplitude of opsin current evoked in L2/3 ACC YFP^+^ interneurons in P15 (Two-way ANOVA p=0.5 in genotype). (G) Representative traces of action potential evoked by a single pulse in L2/3 ACC S5E2^+^ interneurons (4 cycles with 45 s interval). For P15 oIPSC recording, n=52,38 neurons from n=7,6 WT and Shank3B^-/-^ mice. For P15 thalamic input recordings, n=32,35 neurons from n=4,4 WT and Shank3B^-/-^ mice. For adult oIPSC recordings, n=23,25 neurons from n=4,4 WT and Shank3B^-/-^ mice. For AP efficacy recordings, n=19,18 neurons from n=3,3 WT and Shank3B^-/-^ mice. For opsin current recordings, n=19,17 neurons from n=3,3 WT and Shank3B^-/-^ mice. Scatter plots *P<0.05 with two-way ANOVA.

**Supplementary figure 7.**
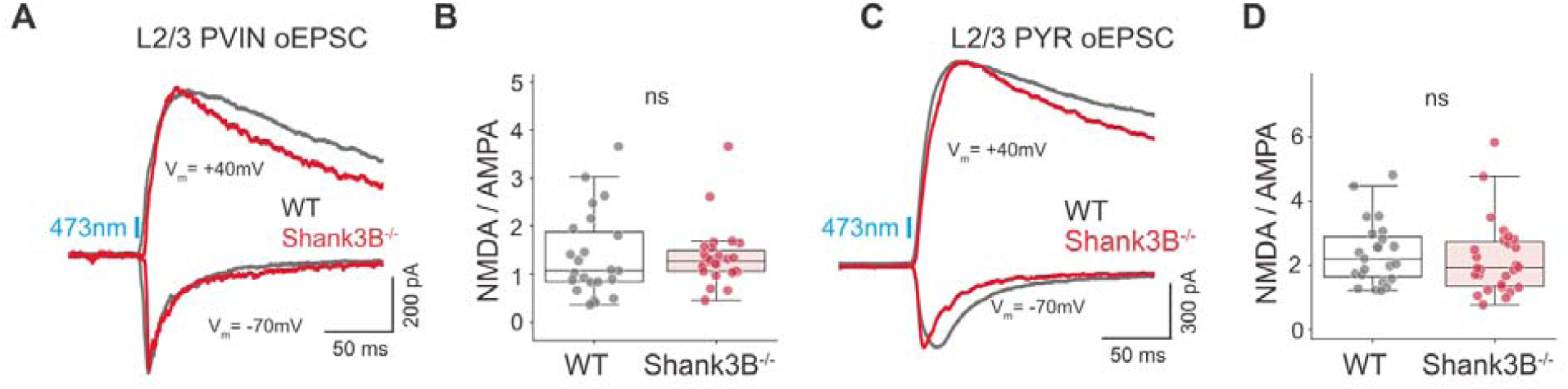
No NMDA / AMPA current ratio difference in both L2/3 PVIN and PYR in ACC of P15 Shank3B^-/-^ mice. **(A)** Representative traces of MD-driven oEPSCs evoked at -70 and +40 mV in L2/3 ACC S5E2^+^ interneurons of WT (black) and Shank3B^-/-^ (red) mice and **(B)** Average of NMDA/AMPA current ratio (Average±SD, WT 1.4±0.9; KO 1.4±0.6, MW test p=0.43). **(C)** Representative traces of MD-driven oEPSCs evoked at -70 and +40 mV in L2/3 ACC PYR of WT (black) and Shank3B^-/-^ (red) mice and **(D)** Average of NMDA/AMPA current ratio (Average±SD, WT 2.4±1.0; KO 2.2±1.1, MW test p=0.37). n=23,27 neurons from n=4,4 WT and Shank3B^-/-^ mice.

**Supplementary figure 8.**
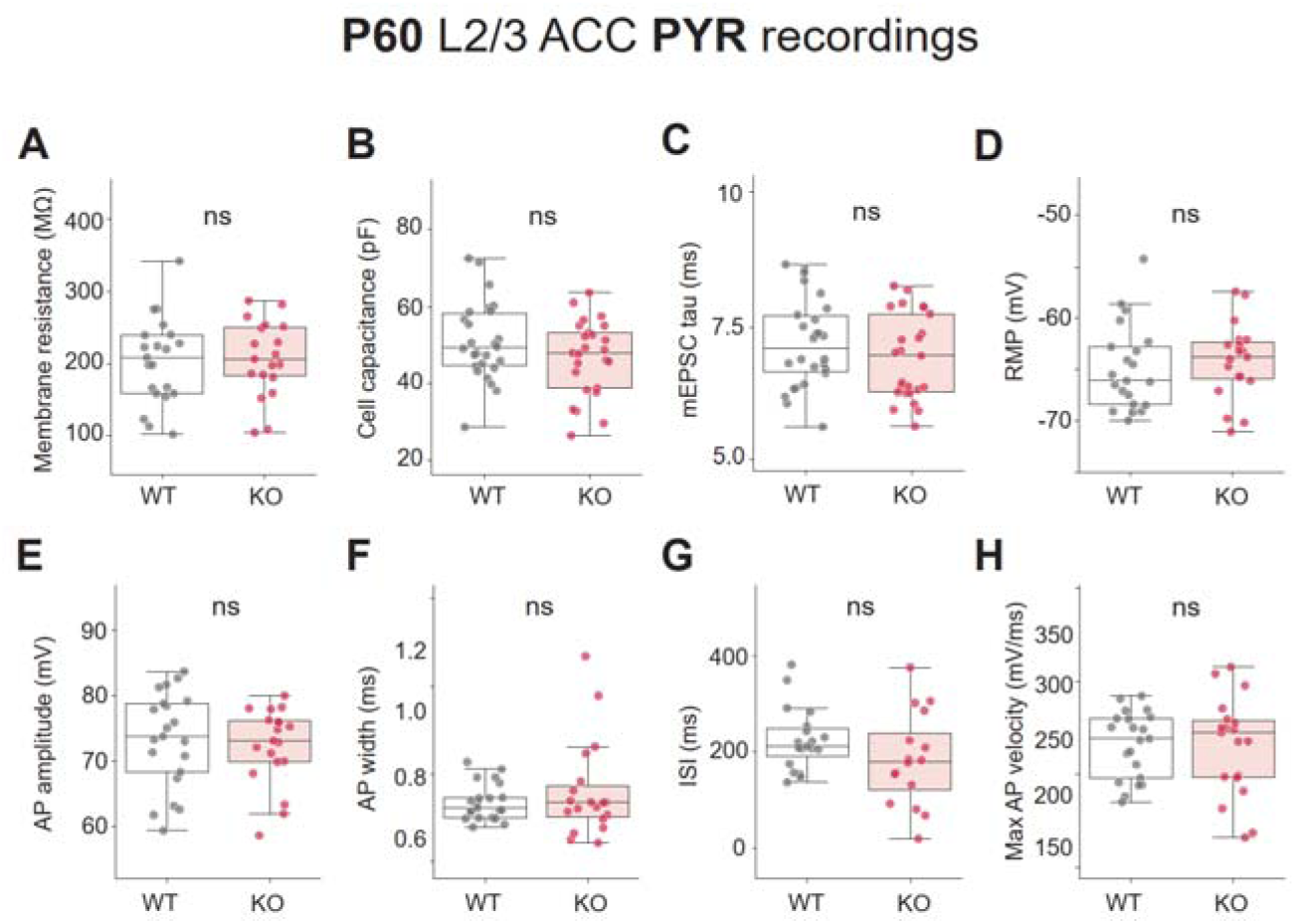
mEPSC and action potential properties of L2/3 PYR in ACC of P60 Shank3B^-/-^ mice. **(A)** Membrane resistance of PYR (Average±SD, WT 215.9±66.4 MΩ; KO 206.7±56.5 MΩ, MW test p=0.77). **(B)** Cell capacitance of PYR (Average±SD, WT 51.0±10.2pF; KO 46.8±9.8pF, MW test p=0.3). **(C)** mEPSC tau of PYR (Average±SD, WT 7.2±0.8 ms; KO 6.9±0.8 ms, MW test p=0.28). **(D)** Resting membrane potential of PYR (Average±SD, WT -64.6±5.0 mV; KO -66.0±4.9 mV, MW test p=0.8). **(E)** AP amplitude PYR (Average±SD, WT 71.6±8.4 mV; KO 72.4±5.9 mV, MW test p=0.95). **(F)** AP width of PYR (Average±SD, WT 0.9±0.1 ms; KO 0.8±0.1 ms, MW test p=0.17). **(G)** Inter-spike interval of PYR (Average±SD, WT 222.6±66.8 ms; KO 175.5±89.8 ms, MW test p=0.08). **(H)** Maximum AP velocity of PYR (Average±SD, WT 223.1±42.8 mV/ms; KO 246.3±41.6 mV/ms, MW test p=0.12). For mEPSC recordings, n=26,25 neurons from n=3,3 WT and Shank3B^-/-^ mice. For current-clamp recordings, n=24,20 neurons from n=3,3 WT and Shank3B^-/-^ mice. Box plots show median and quartile values. Box plots *P<0.05 with MW test.

**Supplementary figure 9.**
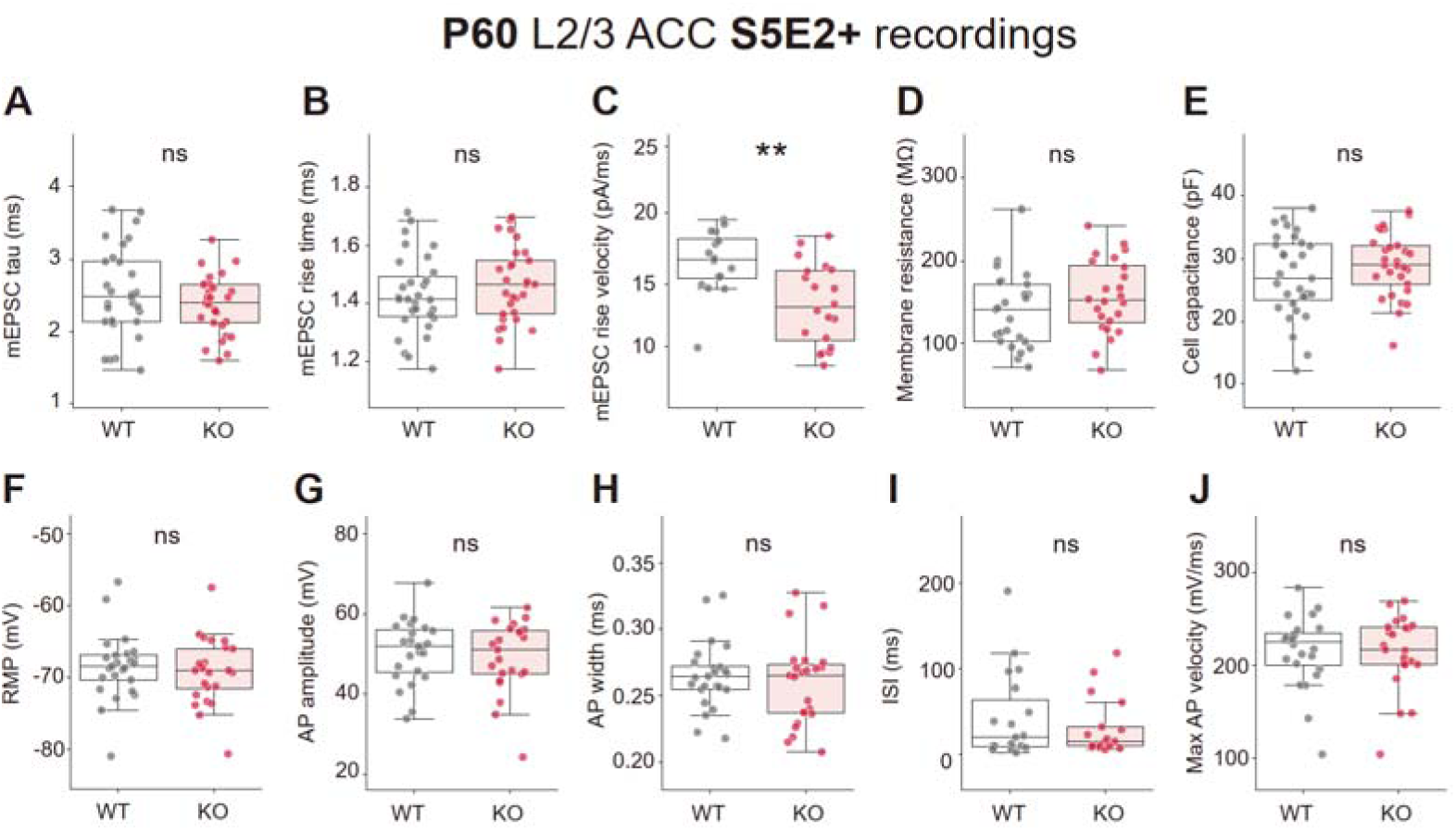
mEPSC and action potential properties of L2/3 S5E2-Tom positive interneurons in ACC of adult Shank3B^-/-^ mice. (A) mEPSC tau of S5E2-Tom+ interneurons (Average±SD, WT 2.5±0.6 ms; KO 2.4±0.4 ms, MW test p=0.4). (B) mEPSC rise time of S5E2-Tom+ interneurons (Average±SD, WT 1.4±0.1 ms; KO 1.5±0.1 ms, MW test p=0.33). (C) mEPSC rise velocity of S5E2-Tom+ interneurons (Average±SD, WT 16.1±2.4 pA/ms; KO 12.9±3.0 pA/ms, MW test p=0.003). (D) Membrane resistance of S5E2-Tom+ interneurons (Average±SD, WT 139.1±45.3MΩ; KO 156.2±44.8MΩ, MW test p=0.16). (E) Cell capacitance of E2-Tom+ interneurons (Average±SD, WT 27.1±6.5 pF; KO 28.9±5.0 pF, MW test p=0.32). (F) Resting membrane potential of S5E2-Tom+ interneurons (Average±SD, WT -68.5±4.7 mV; KO -69.1±4.6 mV, MW test p=0.58). (G) AP amplitude of S5E2-Tom+ interneurons (Average±SD, WT 50.6±7.9 mV; KO 49.2±8.9 mV, MW test p=0.81). (H) AP width of S5E2-Tom+ interneurons (Average±SD, WT 0.3±0.0 ms; KO 0.3±0.0 ms, MW test p=0.6). (I) Inter-spike interval of S5E2-Tom+ interneurons (Average±SD, WT 43.5±51.0 ms; KO 32.1±34.4 ms, MW test p=0.79). (J) Maximum AP velocity of S5E2-Tom+ interneurons (Average±SD, WT 215.5±38.4 mV/ms; KO 214.2±39.4 mV/ms, MW test p = 0.92). For mEPSC recordings, n=30,29 neurons from n=4,4 WT and Shank3B^-/-^ mice. For current-clamp recordings, n=25,24 neurons from n=3,3 WT and Shank3B^-/-^ mice. Box plots show median and quartile values. Box plots **P<0.01 with MW test.

**Supplementary figure 10.**
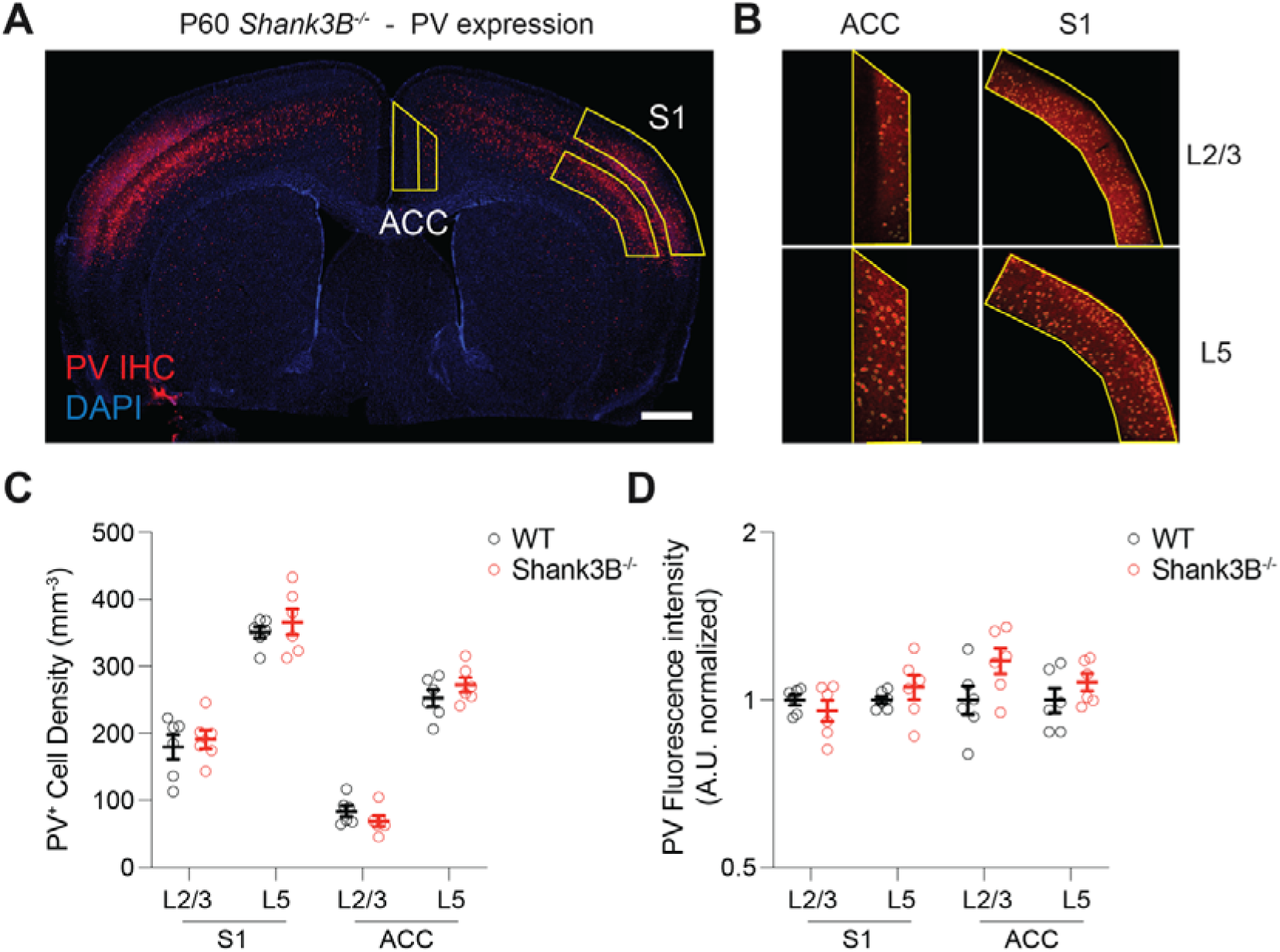
PVIN density and expression intensity in ACC and S1 of adult Shank3B^-/-^ mice. **(A)** Representative coronal section from a P60 Shank3B^-/-^ mouse showing PV immunohistochemistry (red) and DAPI (blue). Regions of interest in the ACC and primary somatosensory cortex (upper lip/forelimb region) were outlined for quantification. Scale bar: 1mm. **(B)** High magnification images of PV+ cells in L2/3 and L5 of ACC and S1 with ROIs used to calculate cell density, illustrating regional and laminar distribution. **(C)** Quantification of PVIN cell density (cells/mm³) for each mouse across cortical layers and regions in WT and Shank3B^-/-^ mice. For each mouse, 4 slices were analyzed and averaged. No significant reduction in PV+ cell density was observed in ACC L2/3 (Average±SD, KO 69.3±19.5; WT 83.6±20.0) and L5 (Average±SD, KO 272.6±27.3; WT 252.6±30.7). S1 layers also showed no significant differences, with PV+ density preserved or mildly elevated in Shank3B^-/-^ mice (Average±SD, S1 L5 KO 366.5±47.4; WT 350.9±21.2; S1 L2/3 KO 191.3±34.1; WT 179.6±44.9). Mixed linear model revealed a significant effect of region (S1 > ACC, p < 0.001) and layer (L5 > L2/3, p < 0.001), but no significant main effect of genotype (p = 0.443) or interaction effects. **(D)** PV fluorescence intensity (arbitrary units, normalized to WT average) was also quantified. Shank3B^-/-^ mice showed significantly lower PV expression in ACC compared to S1 across layers (e.g., ACC L2/3 KO mean ≈ 100.2 ± 15.3 vs. S1 L5 KO ≈ 163.4 ± 11.5). Mixed linear model indicated significant effects of region (p = 0.015) and layer (p = 0.001), with no main effect of genotype (p = 0.561) or interaction terms. N = 6,6 WT and Shank3B^-/-^ male mice. These results indicate no genotype differences in region- and layer-specific PV+ interneuron density or PV expression levels in the ACC and anterior S1 of Shank3B^-/-^ mice.

**Supplementary figure 11.**
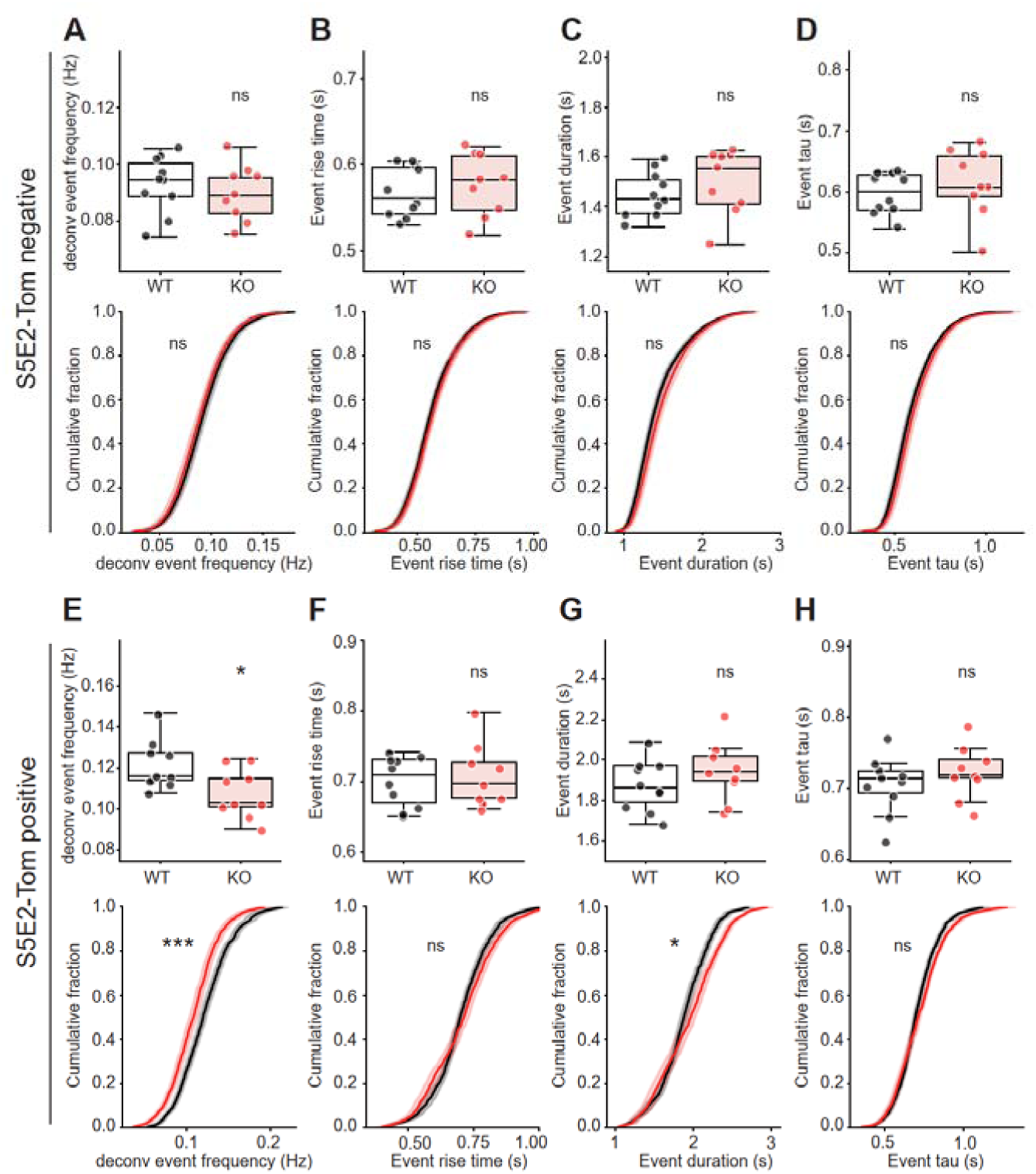
Reduced deconvolved event frequency of S5E2-Tom positive interneurons in dorsal ACC of P16 Shank3B^-/-^ mice was consistent to decrease of z-scored event frequency. (A) Mouse average (upper) and cumulative distribution (lower) of GCaMP8s deconvolved event frequency from S5E2-Tom negative neurons (Average±SD, WT 0.09±0.01Hz; KO 0.09±0.01Hz, MW test p=0.60, KS test p=0.89). (B) Mouse average (upper) and cumulative distribution (lower) of GCaMP8s event rise time from S5E2- Tom negative neurons (Average±SD, WT 0.57±0.03s; KO 0.58±0.04s, MW test p=0.54, KS test p=0.87). (C) Mouse average (upper) and cumulative distribution (lower) of GCaMP8s event duration from S5E2-Tom negative neurons (Average±SD, WT 1.45±0.09s; KO 1.50±0.13s, MW test p=0.21, KS test p=0.29). (D) Mouse average (upper) and cumulative distribution (lower) of GCaMP8s event tau from S5E2-Tom negative neurons (Average±SD, WT 0.6±0.03s; KO 0.61±0.06s, MW test p=0.35, KS test p=0.54). (E) Mouse average (upper) and cumulative distribution (lower) of GCaMP8s deconvolved event frequency from S5E2-Tom positive interneurons (Average±SD, WT 0.12±0.01Hz; KO 0.11±0.01Hz, MW test p=0.03, KS test p=8e-4). (F) Mouse average (upper) and cumulative distribution (lower) of GCaMP8s event rise time from S5E2-Tom positive interneurons (Average±SD, WT 0.70±0.04s; KO 0.71±0.04s, MW test p=0.97, KS test p=0.47). (G) Mouse average (upper) and cumulative distribution (lower) of GCaMP8s event duration from S5E2-Tom positive interneurons (Average±SD, WT 1.88±0.13s; KO 1.95±0.15s, MW test p=0.35, KS test p=0.03). (H) Mouse average (upper) and cumulative distribution (lower) of GCaMP8s event tau from S5E2-Tom positive interneurons (Average±SD, WT 0.71±0.04s; KO 0.72±0.04s, MW test p=0.35, KS test p=0.29). N=10,9 WT and Shank3B^-/-^ mice. Box plots show median and quartile values. Box plots *P<0.05 with MW test. eCDFs *P<0.05, ***P<0.001 with KS test.

**Supplementary figure 12.**
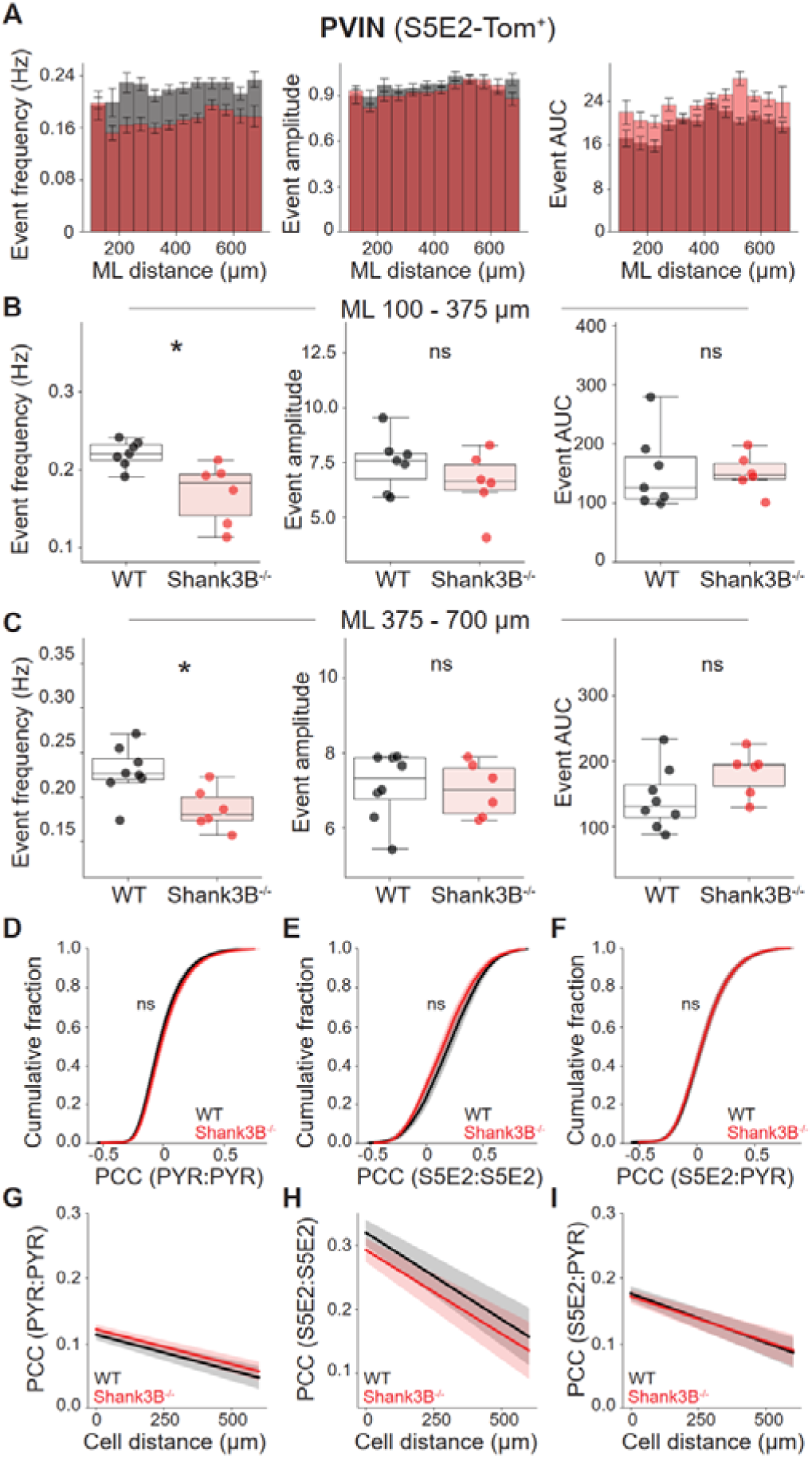
No difference in PVIN activity across cortical layers and normal neural activity correlation in ACC of P15-P17 Shank3B^-/-^ mice. (A) Bar plots of event frequency, amplitude, and AUC of putative PVIN across mediolateral (ML) distance for WT (gray) and KO (red) mice. Cells are grouped in every 50 µm ML distance. (B) Average of GCaMP8s event frequency, amplitude, and AUC of putative PVIN in superficial layer (100<ML≤375 µm), (Average±SD event frequency: WT 0.22±0.02 Hz; KO 0.17±0.04 Hz, MW test p=0.01. Average±SD event amplitude: WT 1.0±0.1; KO 0.9±0.1, MW test p=0.53. Average±SD event AUC, WT 20.3±5.0; KO 20.1±2.5, MW test p=0.73). (C) Average of GCaMP8s event frequency, amplitude, and AUC of putative PVIN in deep cortical layers (ML>375 µm) (Average±SD deep PVIN event frequency: WT 0.23±0.03 Hz; KO 0.19±0.02 Hz, MW test p=0.02. Average±SD deep PVIN event amplitude: WT 1.0±0.1; KO 1.0±0.1, MW test p=0.66. Average±SD deep PVIN event AUC: WT 21.0±4.0; KO 24.3±2.9, MW test p=0.11). (D-F) Cumulative distribution of Pearson correlation coefficient (PCC) of activity between different neuronal populations, PYR-PYR KS test p=0.2. PVIN-PVIN KS test p=1.0. PVIN-PYR KS test p=1.0. (G-I) Linear regressions of PCC against to cell distance. n=8,6 WT and Shank3B^-/-^ mice. Box plots show median and quartile values. Box plots *P<0.05 with MW test. eCDFs *P<0.05 with Kolmogorov-Smirnov test.

**Supplementary figure 13.**
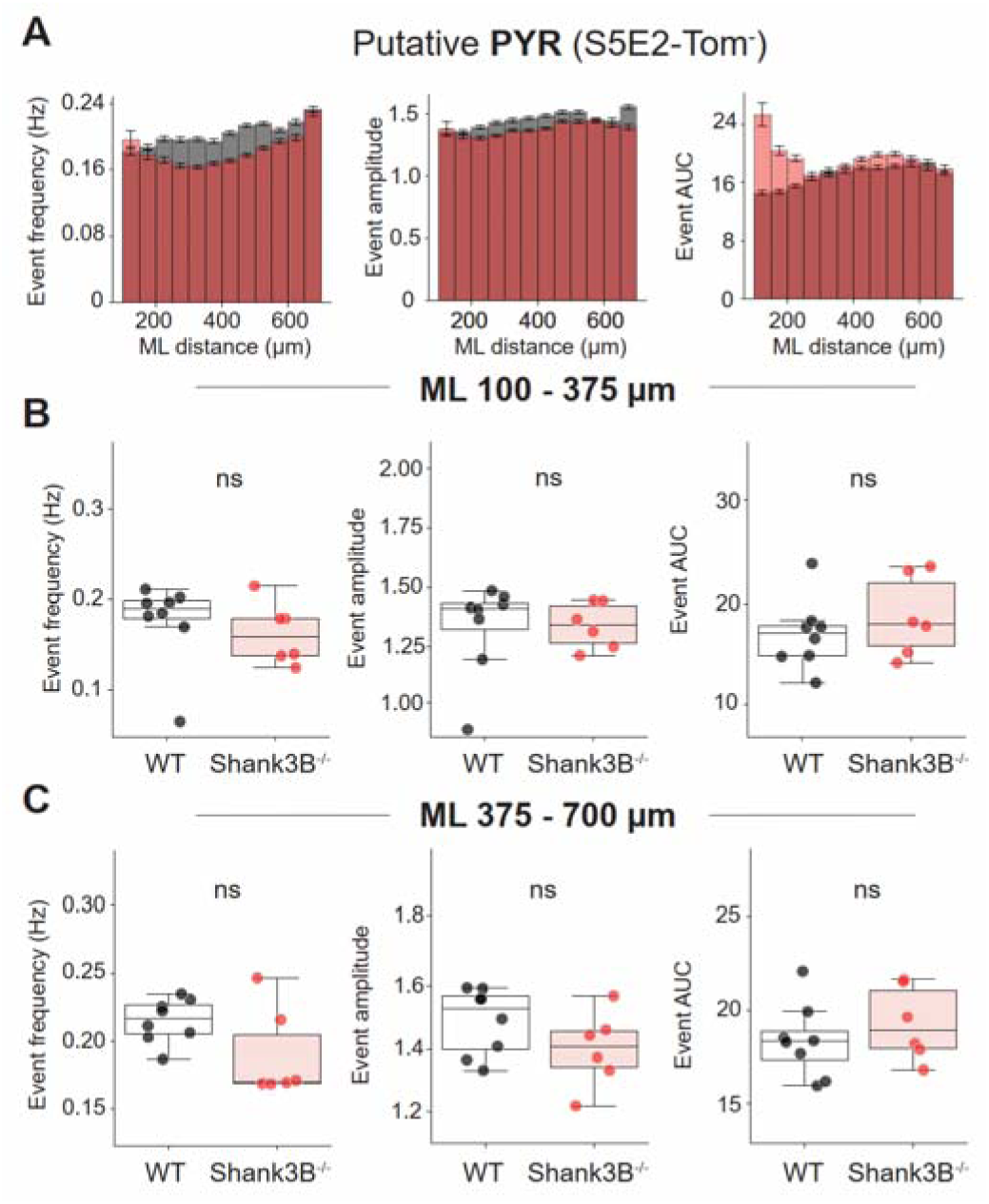
Laminar and mediolateral distance-dependent analysis of calcium event properties in putative pyramidal (S5E2-Tom negative) in Shank3B^-/-^ mice. (A) Bar plots showing the mean ± SEM event frequency, amplitude, and area under the curve (AUC) as a function of mediolateral (ML) distance in putative pyramidal (S5E2-negative) neurons of WT (gray) and KO (red) mice. No significant ML-dependent differences are observed. (B) Box plots showing the event frequency, amplitude, and AUC of calcium events in putative pyramidal neurons in Layer 2/3 ACC of WT and KO mice (Average±SD superficial PYR event frequency: WT 0.18±0.05 Hz; KO 0.16±0.03 Hz, MW test p=0.28. Average±SD superficial PYR event amplitude: WT 1.3±0.2; KO 1.3±0.1, MW test p=0.66. Average±SD superficial PYR event AUC: WT 17.3±3.5; KO 19.1±4.0, MW test p=0.49). (C) Box plots showing the event frequency, amplitude, and AUC of calcium events in putative pyramidal neurons in deep cortical layers of WT and KO mice. (Average±SD deep PYR event frequency: WT 0.22±0.02 Hz; KO 0.19±0.03 Hz, MW test p = 0.14. Average±SD deep PYR event amplitude: WT 1.5±0.1; KO 1.4±0.1, MW test p=0.28. Average±SD deep PYR event AUC: WT 18.2±2.1; KO 19.2±2.1, MW test p=0.66). N=8,6 WT and Shank3B^-/-^ mice. Box plots show median and quartile values. Box plots *P<0.05 with MW test.

## Notes

### Competing Interest Statement

The authors have declared no competing interest.

### Summary of Updates

Added new data and improvements to the manuscript.

